# F-box protein FBXB-65 regulates anterograde transport of UNC-104 through modification near the PH domain

**DOI:** 10.1101/2023.08.13.553108

**Authors:** Vidur Sabharwal, Sri Padma Priya Boyanapalli, Amir Shee, Michael L. Nonet, Amitabha Nandi, Debasish Chaudhuri, Sandhya P. Koushika

## Abstract

Axonal transport is essential for cargo movement between the neuronal cell body and synapses. UNC-104/KIF1A, a Kinesin-3 motor in *C. elegans* that anterogradely transports precursors of synaptic vesicles (pre-SVs), is known to be degraded at synapses through the ubiquitin pathway. Knockdown of the E1 ubiquitin-activating enzyme, *uba-1*, leads to increased accumulation of UNC-104 at neuronal ends and at synapses of touch receptor neurons (TRNs). Loss of the F-box protein FBXB-65, a putative E3 ligase, leads to UNC-104 accumulation at distal ends of neurons, alters net anterograde movement of UNC-104, and the intensity of moving UNC-104 puncta likely bound to cargo without changes in synaptic UNC-104 levels. Using a theoretical model, we analyze the steady state distribution of the anterogradely moving UNC-104 motor. A good agreement between the model and the experimental distributions leads to a crucial hypothesis that UNC-104 may exhibit cooperative binding with moving motor puncta likely associated with cargo, which is regulated by *fbxb-65*. FBXB-65 regulates the modification of UNC-104 motor in a region besides the cargo binding PH-domain. Both *fbxb-65* and UNC-104 motor independent of FBXB-65 regulate the extent of cargo transport in the axon and transport behaviour of cargo at branch points. Our study shows that modification of UNC-104 near its cargo-binding domain may regulate number of motors on the cargo surface and this regulation can fine-tune cargo transport to its destination, the synapse.

## Introduction

Axonal transport is essential for establishing and maintaining neuron structure and function (Maday et al., 2014). The molecular motors kinesin and dynein aid in long-distance axonal transport of cargo synthesized in the neuronal cell body. UNC-104/KIF1A, a kinesin-3 family member, is essential for pre-SV exit from the cell body in multiple model systems (Hall and Hedgecock, 1991; Kumar et al., 2010; Okada and Hirokawa, 1999; Pack-Chung et al., 2007). As axons have a majority of plus-end-out oriented microtubules (Baas et al., 1989; Burton and Paige, 1981; Ghosh-Roy et al., 2012; Heidemann et al., 1981), UNC-104 can drive cargo out of the cell body owing to its ability to walk toward the microtubule plus ends (Okada et al., 1995). Mutations in *C. elegans* UNC-104 that correspond to those found in KIF1A associated with hereditary spastic paraplegia have been shown to hyperactivate UNC-104 by increasing the motors association rate with microtubules, leading to motor depletion from the cell body and its accumulation towards the neuronal distal ends (Chiba et al., 2019; Cong et al., 2021; Niwa et al., 2016). Kinesin-1 has been shown to achieve its characteristic steady-state distribution in neurons based solely on its ability to attach to and dissociate from cargo along with intermittent diffusion (loose-bucket brigade model) (Blasius et al., 2013). Dysregulation of kinesin distribution is often associated with perturbed distribution of its cargo (Chiba et al., 2019; Cong et al., 2021). The altered cargo distribution in turn may lead to neuronal dysfunction. Thus, regulating UNC-104/KIF1A distribution is likely essential for maintaining neuronal homeostasis and preventing the progression of neurological diseases.

Levels of UNC-104 at synapses is shown be regulated by ubiquitination (Kumar et al., 2010), which may also play a role in regulating UNC-104 distribution within neurons. Ubiquitin-mediated degradation also regulates several different kinesins involved in the regulation of cell cycle progression such as Kip1p, CENP-E, Kif26B, and MCAK (Cosper et al., 2023; Gordon and Roof, 2001; Schweiggert et al., 2021; Terabayashi et al., 2012). Many studies have demonstrated that ubiquitin and other ubiquitin-like modifiers that cause post-translational modifications (PTMs) regulate protein function independent of ubiquitin’s role in degrading proteins (Dupré and Haguenauer-Tsapis, 2001; Goo et al., 2015; Govers et al., 1999; Lin et al., 2011; van Delft et al., 1997). Indeed, PTMs such as phosphorylation alter the ability of Kinesin-1 and Kinesin-2 to bind cargo (Guillaud et al., 2008; Liang et al., 2014; Matthies et al., 1993; Yoshimura et al., 2010), and alter the ability of Kinesin-3 to move or bind its adapters (Gan et al., 2020; Hummel and Hoogenraad, 2021; Kevenaar et al., 2016). PTMs on motors or motor adapters may also regulate the ability of a kinesin to switch between inactive and active conformations (Cai et al., 2007; Hammond et al., 2009). Whether ubiquitination or other PTMs modify kinesins to help maintain their distribution remains poorly understood (Hong et al., 2018).

Both ubiquitin and ubiquitin-like modifiers are added to lysine residues on protein substrates (Mattiroli and Sixma, 2014). Ubiquitination and its related ubiquitin-like modifications use E1, E2, and E3 sequentially to achieve substrate specificity (Nagy and Dikic, 2010). In *C. elegans*, *uba-1* is the sole E1 ubiquitin-activating enzyme and thus regulates all ubiquitination within the animal (Kulkarni and Smith, 2008). Attachment of ubiquitin or ubiquitin-like modifications can mutually depend on each other via: (i) using the same family of E3 ligases to attach different modifiers (Chu and Yang, 2011; Oved et al., 2006; Schmidt and Dikic, 2005), (ii) attach an ubiquitin-like modifier to an ubiquitin E3 ligase to control its activity (Liu and Xirodimas, 2010; Ohh et al., 2002; Ohki et al., 2009; Oved et al., 2006; Tatham et al., 2008; Uzunova et al., 2007), and (iii) attach a ubiquitin-like modifier to the substrate altering its ability to be ubiquitinated (Sun et al., 2007; Xie et al., 2007). Additionally, the same protein substrate can be regulated by multiple E3s that target different sites on the substrate (Haglund et al., 2003). In summary, the interplay between ubiquitin and ubiquitin-like modifiers can result in complex regulation of substrate modification that may lead to different outcomes of the substrate.

We have previously shown that UNC-104 is degraded in a ubiquitin-dependent manner in *C. elegans* neurons (Kumar et al., 2010). In our current study, using a combination of *in vivo* experiments and analytical theory, we show that a putative E3 ligase, *fbxb-65,* regulates UNC-104 distribution in *C. elegans* neurons. Our study suggests that *fbxb-65* also regulates UNC-104 levels on the cargo surface, potentially by regulating modification of UNC-104 close to the cargo-binding Pleckstrin Homology (PH) domain in the 1386-1421 amino acid (aa) region. Analysis of the UNC-104 motile cluster distributions using our theoretical expression, suggests cooperative binding of motor proteins on the cargo surface, and modifications of the binding rates under FBXB-65 regulation. Lack of FBXB-65 or the region it modifies in UNC-104 regulates the distribution of pre-SVs, movement of pre-SVs along the axon and at branch points in axons. Therefore, FBXB-65*-*mediated modification of UNC-104 likely maintains the distribution of the motor and its cargo RAB-3 by reducing the total UNC-104 bound to cargo.

## Results

### I. *fbxb-65* regulates distal end accumulation of UNC-104

UNC-104 and its mammalian orthologue KIF1A are likely ubiquitinated (Kumar et al., 2010; Ordureau et al., 2020; Oshikawa et al., 2012; Sarraf et al., 2013; Vogl et al., 2020) and rapidly turned over (Cohen et al., 2013; Fornasiero et al., 2018; Huang et al., 2020; Mathieson et al., 2018). Previously, a temperature-sensitive strain of *uba-1(it129ts)* grown at a restrictive temperature has been reported to show UNC-104::GFP accumulation specifically at synapses (Kumar et al., 2010). Here we describe a putative E3 ligase, *fbxb-65*, knockdown of which phenocopies UNC-104 distribution changes similar to that seen in *uba-1* RNAi animals.

The TRN-specific knockdown of *uba-1* results in an increase in TRN-expressed UNC-104::GFP levels in ~90% animals at the PLM distal end compared to that in Empty Vector RNAi (hereon referred to as control) animals [Fig. 1A]. *fbxb-65* RNAi shows distal end accumulation of UNC-104::GFP in ~62% of animals compared to that seen in controls [Fig. 1A]. In control animals, UNC-104::GFP shows a steep increase in intensity at the last 5 μm before the distal end [Fig. S1B]. In contrast, in *uba-1* RNAi and *fbxb-65* RNAi animals, UNC-104::GFP intensity increases in the last 50 μm of the PLM process, with the greatest accumulation at neuronal distal tips [Fig. S1B]. Additionally, TRN-specific expression of FBXB-65::GFP shows diffuse expression throughout the neuron [Fig. S1A]. Thus, FBXB-65 can potentially influence UNC-104 across the entire neuron thereby controlling UNC-104 steady-state distribution.

**Figure 1.**
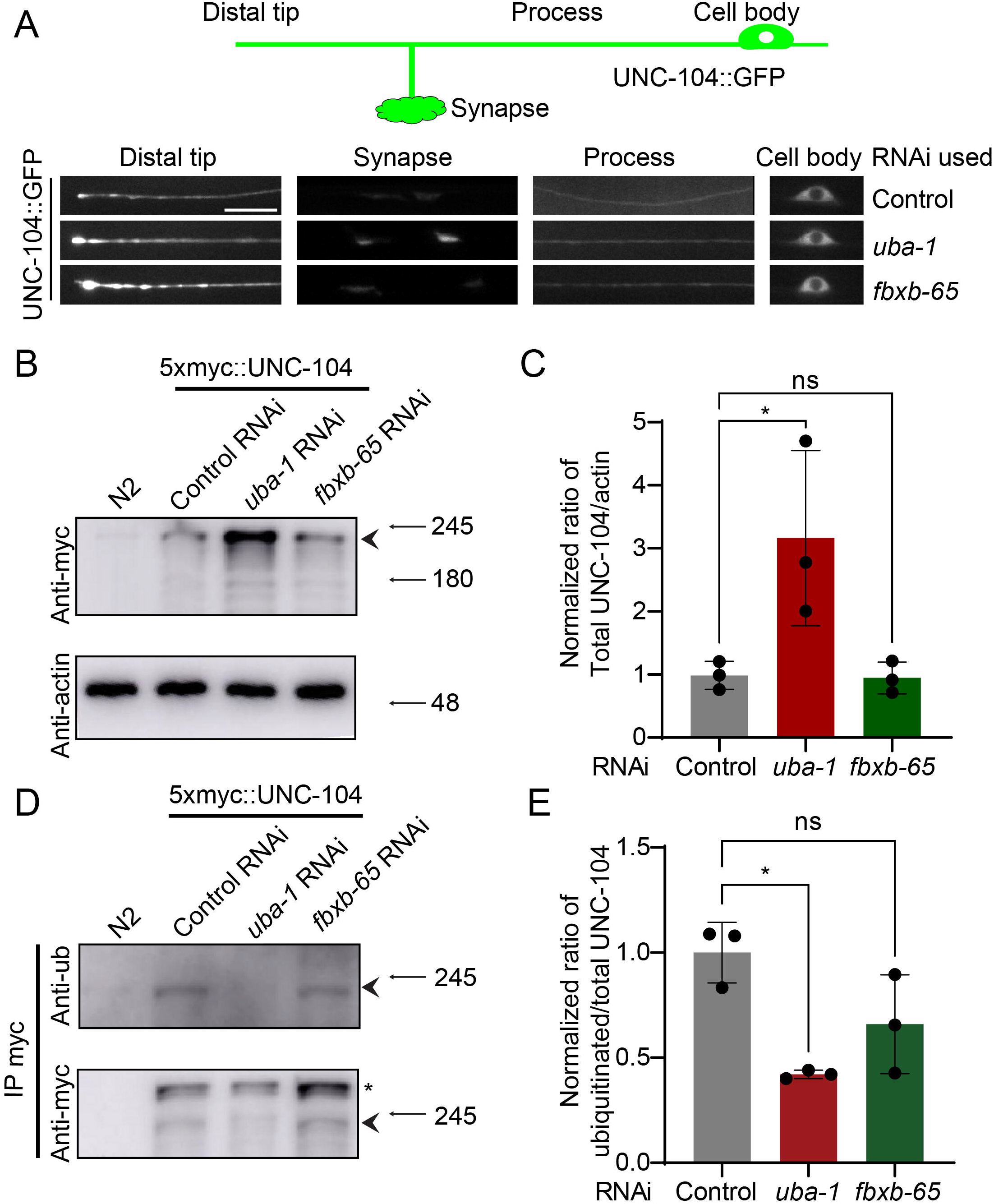
UNC-104 distribution is regulated by *uba-1* and *fbxb-65*. A) Schematic representing the regions in the PLM neuron that were imaged along with representative images of UNC-104::GFP intensity at these regions, namely the cell body, proximal process (70 µm from the cell body), and the distal end in the strain TT2440 treated with either control, *uba-1*, or *fbxb-65* RNAi. Scale bar 10 µm. B) Western blot analyses either from whole worm N2 or TT3185 lysates treated with either control, *uba-1*, or *fbxb-65* RNAi. β-Actin served as a loading control. The arrow head indicates the UNC-104 band used for intensity quantitation. Molecular weights (in kDa) marked with arrows. C) Bar plot of quantitation of total UNC-104 band intensity normalized to the actin loading control. Results are plotted as Mean ± SD (N=3 biological repeats represented as filled circles). *p<0.05 (One-way ANOVA with Dunnett’s multiple comparisons test). D) Western blot analyses from total UNC-104 normalized IP enriched fractions of TT3185 treated with either control, *uba-1*, or *fbxb-65* RNAi probed with anti-ubiquitin (FK2) followed by stripping and re-probing with anti-myc (AE010). * indicates a non-specific band much higher than 250 kDa observed post-IP. The arrow head indicates the UNC-104 band used for intensity quantitation. Molecular weights (in kDa) marked with arrows. E) Bar plot of quantitation of ubiquitinated UNC-104 intensity normalized by the total UNC-104 intensity. Results are plotted as Mean ± SD (N=3 biological repeats represented as filled circles). *p<0.05 (One-way ANOVA with Dunnett’s multiple comparisons test)

RNAi of *uba-1* or *fbxb-65* does not significantly alter UNC-104 RNA levels [Fig. S1C, Table S12], suggesting that knocking down these genes likely regulates UNC-104 distribution through post-translational processes. UNC-104::GFP levels increase at the PLM synapse only in *uba-1* RNAi but not substantially in *fbxb-65* RNAi animals [Fig. 1A, Fig. S1D, Table S12]. Pan-neuronal knockdown of *uba-1* leads to a 3-fold increase in total UNC-104 protein levels, while a similar knockdown of *fbxb-65* does not alter UNC-104 protein levels [Fig. 1B,C, Table S4]. Together these data suggest *fbxb-65* does not regulate UNC-104 protein levels in contrast to that seen in *uba-1*.

UBA-1 specifically catalyzes ubiquitin activation in *C. elegans* (Kulkarni and Smith, 2008). However, E3 ligases like FBXB-65 may enable modification by other ubiquitin-like modifiers as well (Kanei-Ishii et al., 2012). Hence, using a pan anti-ubiquitin antibody on immunoprecipitated UNC-104, we assessed if the ubiquitinated fraction of UNC-104 is altered upon RNAi-mediated knockdown of *uba-1* and *fbxb-65*. Only *uba-1* RNAi shows a significant reduction (58%) while *fbxb-65* RNAi shows a non-significant reduction (34%) in ubiquitinated UNC-104 levels compared to the control [Fig. 1D,E, Table S4]. The larger reduction seen in *uba-1* RNAi compared to *fbxb-65* RNAi may indicate multiple ubiquitination pathways that may modify UNC-104.

The above findings indicate that *fbxb-65* knockdown shows only a subset of *uba-1* RNAi phenotypes, namely UNC-104 accumulation at the distal ends; hence, *fbxb-65* may predominantly regulate UNC-104 along the neuronal process rather than at synapses.

### II. *fbxb-65* knockdown alters the number and intensity of moving UNC-104 puncta

Single motor movement properties can change the overall distribution of the motor along neuronal processes (Chiba et al., 2019; Cong et al., 2021). To assess the movement of the motor we examined movement of UNC-104::GFP in the TRNs (Klopfenstein et al., 2002; Kumar et al., 2010; Wu et al., 2016; Zhou et al., 2001).

Using kymographs of our movies of RNAi-treated *C. elegans*, we measured the flux, velocity, and net displacement of UNC-104::GFP particles [Fig. 2A]. Most of the UNC-104::GFP signal seems diffuse, with intermittent visible moving puncta [Movie 1]. The average net displacement (~7 μm), the fraction of UNC-104::GFP trajectories with run length >7 μm, and velocity (~1 μm s^-1^) are not significantly different between control and *fbxb-65* RNAi animals [Figs. 2B,C, S2B, Tables S5, S13]. There is a 43% increase in UNC-104::GFP anterograde flux, and a ~140% increase in UNC-104::GFP retrograde flux in *fbxb-65* RNAi compared to that in control RNAi [Fig. 2D,E, Table S5]. Additionally, the intensity of moving UNC-104::GFP particles is ~148% greater in *fbxb-65* RNAi than in the control [Fig. 2F, Table S5, Movies 1, 2].

**Figure 2.**
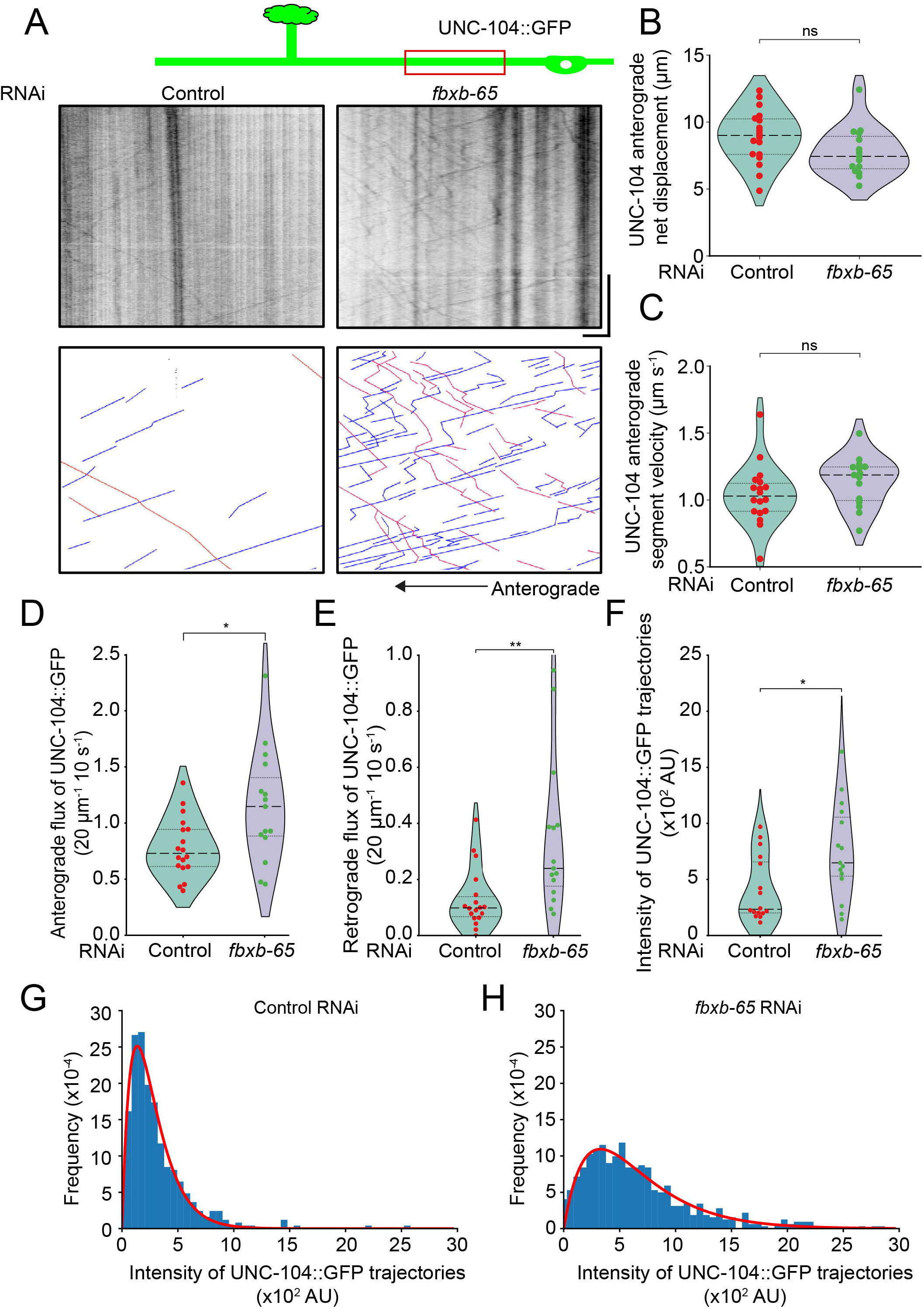
*fbxb-65* RNAi increases flux and intensity of UNC-104 single particle tracks compared to control. A) Representative kymographs for UNC-104::GFP 70 µm from the cell body in the strain TT2440 treated with either control or *fbxb-65* RNAi along with their respective manually-traced annotations shown at the bottom. Blue lines represent a net anterograde motion, while red lines represent a net retrograde motion. Scale x-axis 10 µm, y-axis 30 s. B) Net displacement of anterogradely-moving UNC-104::GFP trajectories assessed from the strain TT2440 treated with either control (N=18) or *fbxb-65* (N=15) RNAi. Data are represented as violin plots with individual data points along with the median (dashed line), 25^th^ and 75^th^ percentile marked (dotted lines); ns-non-significant (Unpaired Student’s *t*-test two-tailed). C) Velocities of uninterrupted anterogradely-moving UNC-104::GFP trajectories assessed from the strain TT2440 treated with either control (N=18) or *fbxb-65* (N=15) RNAi. Data are represented as violin plots with individual data points along with the median (dashed line), 25^th^ and 75^th^ percentile marked (dotted lines). ns-non-significant (Unpaired Student’s *t*-test two-tailed). D) Number of anterograde UNC-104::GFP trajectories normalized to a 20 µm and 10 s region of the kymograph assessed from the strain TT2440 treated with either control (N=18) or *fbxb-65* (N=15) RNAi. Data are represented as violin plots with individual data points along with the median (dashed line), 25^th^ and 75^th^ percentile marked (dotted lines). *p<0.05 (Unpaired Student’s *t*-test two-tailed). E) Number of retrograde UNC-104::GFP trajectories normalized to a 20 µm and 10 s region of the kymograph assessed from the strain TT2440 treated with either control (N=18) or *fbxb-65* (N=15) RNAi. Data are represented as violin plots with individual data points along with the median (dashed line), 25^th^ and 75^th^ percentile marked (dotted lines). **p<0.01 (Mann–Whitney U test two-tailed). F) Background subtracted intensity of anterograde UNC-104::GFP trajectories assessed from the strain TT2440 treated with either control (N=18) or *fbxb-65* (N=15) RNAi. Data are represented as violin plots with individual data points along with the median (dashed line), 25^th^ and 75^th^ percentile marked (dotted lines). *p<0.05 (Mann–Whitney U test two-tailed). G) Frequency distribution of background subtracted UNC-104 intensity averaged over an entire individual run assessed from the strain TT2440 treated with control (N=18) RNAi. The distribution is overlaid with a probability distribution function (red line) generated from our theoretical equation (6) assuming γ=1 and *μ* = 7.33 × 10^-3^. Parameters were derived from the datasets by least square fitting. G) Frequency distribution of background subtracted UNC-104 intensity averaged over an entire individual run assessed from the strain TT2440 treated with *fbxb-65* (N=15) RNAi. The distribution is overlaid with a probability distribution function (red line) generated from our theoretical equation (6) assuming γ=1 and *μ* = 3.05 × 10^-3^. Parameters were derived from the datasets by least square fitting.

Thus, knocking down *fbxb-65* leads to an increase in the intensity of moving UNC-104::GFP trajectories in addition to more moving UNC-104::GFP particles. These moving UNC-104::GFP likely reflect motors bound to its cargo. Since *fbxb-65* RNAi does not alter the levels of total UNC-104 or synaptic UNC-104::GFP, *fbxb-65* may regulate UNC-104 largely within the neuronal process.

### III. *fbxb-65* knockdown increases UNC-104 accumulation at cut sites post-axotomy

*fbxb-65* RNAi led to an increase in flux of UNC-104. Net movement of motors along the axon can also be assessed by an injury/axotomy-based assay (Abay et al., 2017; Kumar et al., 2010; Li et al., 1999; Okada and Hirokawa, 1999; Rao et al., 2008). It has been reported that in *uba-1* mutants there was increased accumulation of the motor at the distal cut site compared to wild type animals (Kumar et al., 2010), suggesting an increase in retrograde movement of UNC-104 at longer time scales in *uba-1*. Therefore, to assess if this assay also showed increased UNC-104 movement in *fbxb-65* RNAi, we assessed accumulation of UNC-104 in the first 120 s after injury as well as 1 h post-injury.

We used a femtosecond laser to cut the neuronal process ~100 μm away from the cell body and accumulation of UNC-104::GFP was tracked at the cut sites proximal and distal to the cell body in the first 120 s post-axotomy [Fig. 3A,B]. In *fbxb-65* RNAi animals, the accumulation of UNC-104::GFP at the proximal cut site is 1.5-fold faster and 40% higher compared to the control [Fig. 3C,D,E, Table S6]. The UNC-104::GFP intensity at the distal cut site in *fbxb-65* RNAi animals remains unchanged, which contrasts with that in the control, where UNC-104::GFP first increases and then decays post-injury [Fig. S2A]. At 1 h post-injury, UNC-104::GFP intensity at the proximal cut site continues to remain high, with *fbxb-65* RNAi animals showing a 75% higher intensity than the control [Fig. 3F,G], while the distal cut site shows a 70% increase in intensity [Fig. 3H, Table S6]. Thus, *fbxb-65* knockdown likely increases both anterograde and retrograde movement of UNC-104.

**Figure 3.**
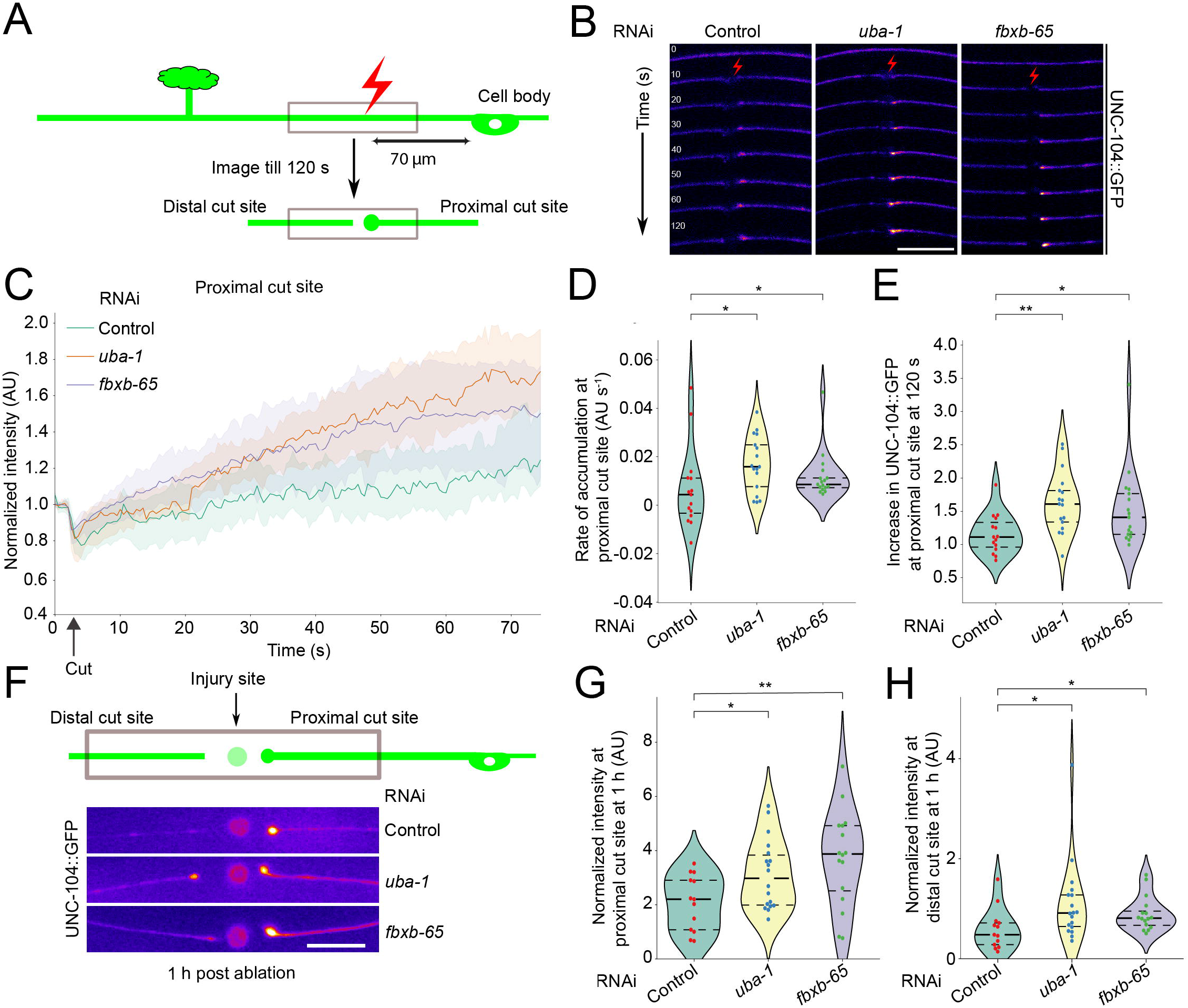
UNC-104::GFP proximal accumulation post-injury is enhanced in *fbxb-65* and *uba-1* RNAi. A) Schematic showing the injury paradigm with LASER-induced ablation around 70 µm from the cell body. B) Representative images of UNC-104::GFP before and after ablation in the strain TT2440 treated with either control, *uba-1*, or *fbxb-65* RNAi at time points 0 s (before ablation) up to 120 s after ablation. Scale bar 10 µm. C) UNC-104::GFP intensity normalized to the average intensity of 5 frames pre-ablation plotted for a region including 5 µm of the cut site proximal to the cell body in the strain TT2440 treated with either control (N=15), *uba-1* (N=17), or *fbxb-65* (N=17) RNAi. The solid line represents the mean and the shaded region represents the 95% confidence interval for ablation aligned (labeled as cut) time series intensities of n>15 for all conditions. D) Rate of accumulation plotted for UNC-104::GFP intensity in the 5 µm region of the cut site proximal to the cell body in the strain TT2440 treated with either control (N=15), *uba-1* (N=17), or *fbxb-65* (N=17) RNAi starting from 20 s till 40 s. Data are represented as violin plots with individual data points along with the median, 25^th^ and 75^th^ percentile marked. *p<0.05 (Mann–Whitney–Wilcoxon test with Bonferroni correction). E) UNC-104::GFP intensity at 120 s normalized to pre-ablation intensity plotted in the 5 µm region of the cut site proximal to the cell body in the strain TT2440 treated with either control (N=15), *uba-1* (N=17), or *fbxb-65* (N=17) RNAi. Data are represented as violin plots with individual data points along with the median, 25^th^ and 75^th^ percentile marked. *p<0.05 **p<0.01 (Mann–Whitney–Wilcoxon test with Bonferroni correction). F) Schematic and representative images of UNC-104::GFP in worms rescued post ablation and imaged 1 h later. The strain used was TT2440 treated with either control, *uba-1*, or *fbxb-65* RNAi. G) UNC-104::GFP intensity in a 5 µm region at the proximal cut site normalized to intensity in a 5 µm region in the proximal neuronal process 20 µm away from the cut site in the strain TT2440 treated with either control (N=13), *uba-1* (N=18), or *fbxb-65* (N=16) RNAi. Data are represented as violin plots with individual data points along with the median, 25^th^ and 75^th^ percentile marked. *p<0.05 **p<0.01 (Mann–Whitney–Wilcoxon test with Bonferroni correction). H) UNC-104::GFP intensity in a 5 µm region at the distal cut site normalized to intensity in a 5 µm region in the distal neuronal process 20 µm away from the cut site in the strain TT2440 treated with either control (N=13), *uba-1* (N=18), or *fbxb-65* (N=16) RNAi. Data are represented as violin plots with individual data points along with the median, 25^th^ and 75^th^ percentile marked. *p<0.05 (Mann–Whitney–Wilcoxon test with Bonferroni correction).

At 1 h post-injury, *uba-1* RNAi leads to a greater accumulation of UNC-104::GFP, with an increase of 95% and 35% at the distal and proximal cut sites, respectively, compared to that in the control [Fig. 3F,G,H, Table S6] (Kumar et al., 2010). At 120 s post-injury, there is a 2-fold faster and 50% higher accumulation of UNC-104::GFP in the proximal cut site in *uba-1* RNAi animals compared to that in the control [Fig. 3C,D,E, Table S6]. Immediately post-injury, UNC-104::GFP intensity at the distal cut site reduces in *uba-1* RNAi animals [Fig. S2A]. Thus, *uba-1* knockdown likely increases UNC-104’s anterograde movement at shorter time scales but may lead to a larger net retrograde movement at longer time scales.

The UNC-104::GFP accumulation at the proximal cut site immediately post-injury is increased in both *fbxb-65* and *uba-1* RNAi animals compared to the control. At 1 h post axotomy in *fbxb-65* RNAi animals, UNC-104::GFP accumulation at the proximal and distal cut sites increases to a similar extent. In contrast, *uba-1* RNAi leads to increased accumulation at the distal end. Therefore, *fbxb-65* may regulate overall UNC-104 anterograde and retrograde flux within the neuronal process. *uba-1* increases UNC-104 retrograde flux [Fig.], possibly due to a decrease in UNC-104 degradation at synapses, as previously suggested (Kumar et al., 2010). The increased accumulation of UNC-104 at the neuronal ends of the TRNs may arise from altered net movement of the motors in *fbxb-65* knockdown.

### IV. Ubiquitin-like modifiers may regulate binding of UNC-104 to cargo

To test if ubiquitination-induced alteration in UNC-104::GFP attachment and detachment in the motor puncta trajectories can be distinguished, we first present a theoretical analysis considering the time evolution of the probability distribution *P*(*n,t*) that a trajectory has *n* motors engaged in its transport on microtubules. This evolution can be expressed in terms of loss *u*_−_(*n*) and gain *u*_+_(*n*) rates of binding and unbinding of motors (see Materials and Methods for the full theory). A punctum cannot lose motors unless it has a motor already present, thus the minimal assumption of the loss rate is

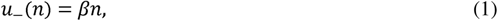

up to linear order in *n*. Here *β* is the detachment rate constant. Similarly, within the same linear approximation, the gain rate can be expressed as

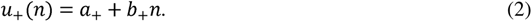

The first term *a*_+_ in the above expression [Eqn. 2] denotes a constant binding rate while the second term *b*_+_*n* denotes a cooperative binding rate. In the presence of the second term, the probability of binding new motor proteins increases with the number of existing motor proteins in a punctum. With these forms for the loss and gain rates [Eqns. 1, 2], the steady state distribution of motor proteins in a punctum is given by (see Materials and Methods),

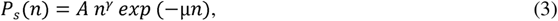

where *γ a*_+_/*β* − 1 and *μ* = 1 − *b*_+_/*β*. Fitting Eqn. 3 to the experimentally obtained distributions [Fig. 2G,H], it becomes clear that the assumption of cooperative binding (*b*_+_ ≠ 0) is indeed essential to explain the observed steady state distributions of the control and *fbxb-65* RNAi. Without cooperative binding (*b*_+_ = 0), the distribution [Eqn. 3] takes the form *P_S_*(*n*) = *A n^γ^ exp* (-*n*) which fails to capture the difference in the mode 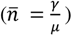 and variance (*σ*^2^*_n_* = 2/μ^2^, see Materials and Methods) of the distributions between control and *fbxb-65* RNAi.

The above expression of *P_S_*(*n*) [Eqn. 3] fits well with the probabilities obtained from the intensity distributions of moving UNC-104::GFP puncta for the control [Fig. 2G] and *fbxb-65* RNAi [Fig. 2H] with *γ* = 1 and *μ_control_* > *μ_fbxb-65_* [Table S17]. Since *μ_control_* > *μ_fbxb-65_*_’_ *b*_+_ is smaller in the control and larger in *fbxb-65* RNAi, suggesting that *fbxb-65* knockdown can increase UNC-104’s ability to bind cargo to facilitate movement. Further, since *γ* = 1 fits most of our data well, it is parsimonious that ubiquitination regulates the cooperative recruitment of UNC-104 on the cargo surface and likely does not alter UNC-104’s detachment.

### V. UNC-104 is likely modified near its PH domain

UNC-104 contains multiple lysines that can be used to attach ubiquitin or other ubiquitin-like modifiers. Since *fbxb-65* does not appear to regulate overall ubiquitination levels of UNC-104, this gene may potentially alter a single mono-ubiquitin addition or indirectly affect attachment of other ubiquitin-like modifiers. The addition of ubiquitin or ubiquitin-like modifiers will change UNC-104 mobility only by ~8 kDa, making it difficult to identify the change on a western blot since UNC-104 is ~180 kDa. Thus, we created multiple fragments of UNC-104 spanning the entire coding region of UNC-104 to identify the region that demonstrates an *fbxb-65*-dependent increase in molecular weight.

We created four different fragments, encompassing the motor domain (fragment 1), the coiled-coil domain important for UNC-104 dimerization (fragment 2), the stalk domain (fragment 3), and the cargo-binding PH domain (fragment 4) [Fig. 4A]. Upon expression, the construct containing fragment 4 exhibits two distinct bands separated by ~8 kDa [Fig. 4B], which is the size expected for mono-ubiquitin or mono-ubiquitin-like modifiers (e.g., SUMO, NEDD, URM, and UFM). Furthermore, the relative ratio of the higher to lower molecular weight PH fragments significantly reduces in pan-neural *fbxb-65* RNAi [Fig. 4C,D], suggesting that *fbxb-65* may regulate modification of UNC-104 close to its cargo-binding PH domain. We also attempted to assess if the top band of fragment 4 is ubiquitinated using anti-ubiquitin antibodies; however, no signal was obtained for the observed size [Fig. S3C]. Since the FBXB-65 dependent higher molecular weight band, is not recognized by anti-ubiquitin antibodies, and because *fbxb-65* knockdown does not significantly change overall levels of ubiquitinated UNC-104 [Fig. 1D,E] the modification may be ubiquitin-like and depend on other ubiquitin-like E1s. *C. elegans* has five E1 activating enzymes, one each for ubiquitin, SUMO, NEDD8, URM1, and UFM1, encoded by genes *uba-1*, *uba-2*, *rfl-1*, *moc-3*, and *uba-5*, respectively. Pan-neuronal knockdown of these E1 genes does not show a significant reduction in the modification of fragment 4, except for *uba-1* RNAi, which shows a 20% reduction [Fig. S3A,B, Table S14]. We could not confirm the nature of the modification on fragment 4 due to the lack of antibodies that recognize the *C. elegans* ubiquitin-like modifiers.

**Figure 4.**
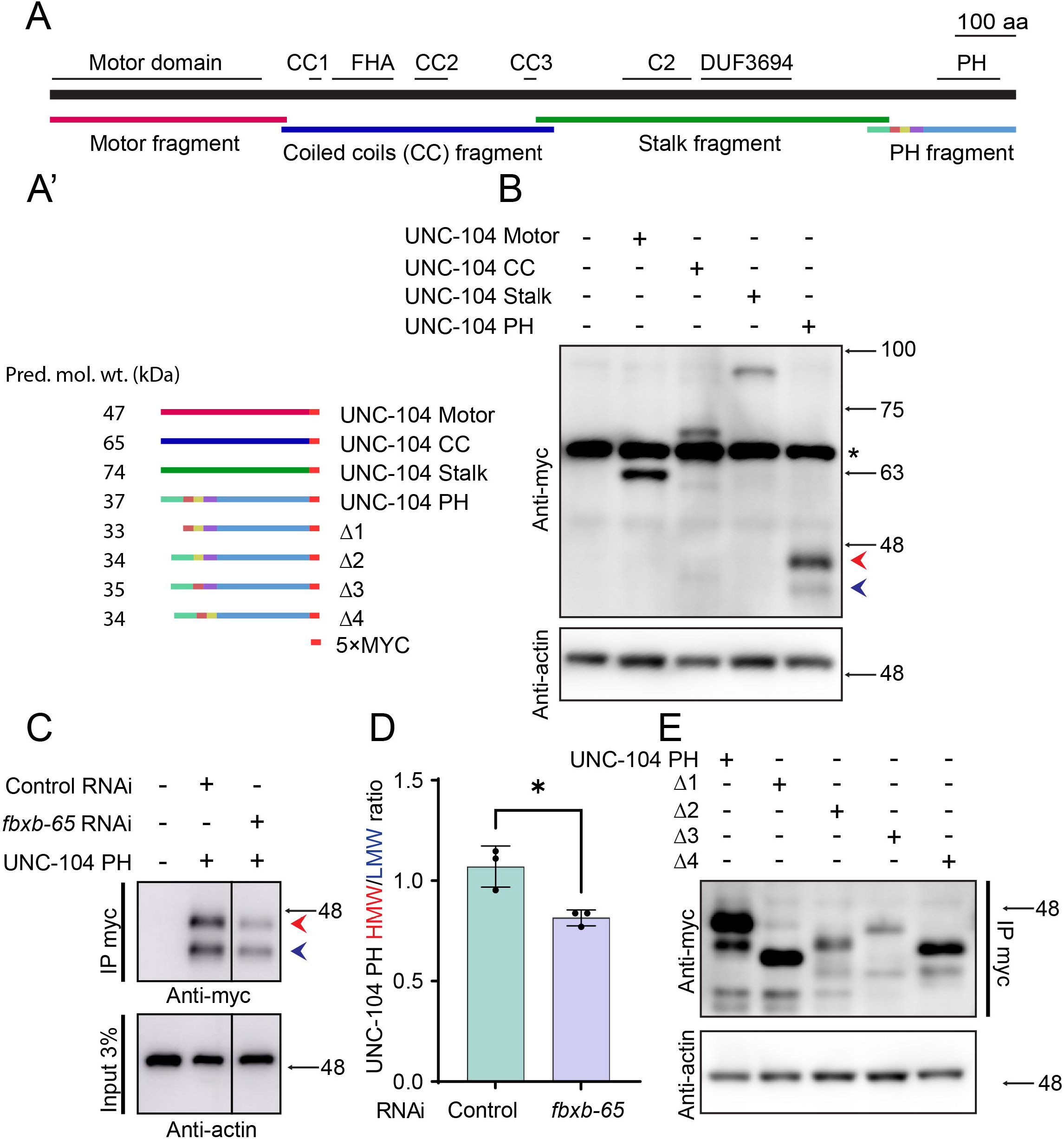
A region near UNC-104 PH domain is likely modified by *fbxb-65*. A) Illustration of known UNC-104 domains marked on top and the different fragments created on the bottom of the dark thick line representing the UNC-104 sequence. Scale 100 amino acids. A’ lists the different combinations of fragments fused with a 5×MYC tag with their predicted molecular weights mentioned. B) Western blot analyses from whole worm lysates expressing fragments of UNC-104 (Motor, CC, Stalk, and PH domain-containing fragment) fused with a 5×MYC tag probed with anti-myc. The blots were stripped and re-probed with anti-β-Actin to serve as a loading control. * indicates a non-specific band just above 63 kDa. The two bands of UNC-104 PH::5×MYC are marked with arrow heads as HMW (High Molecular Weight) in red and LMW (Low Molecular Weight) in blue. Molecular weights (in kDa) are marked with arrows. C) Western blot analyses from anti-myc IP enriched fractions of TT3158 treated with either control or *fbxb-65* RNAi. The input served as a loading control probed with β-Actin. The two bands of UNC-104 PH::5xMYC are marked with arrow heads as HMW (High Molecular Weight) in red and LMW (Low Molecular Weight) in blue. Molecular weights (in kDa) are marked with arrows. D) Bar plot representing mean intensity of the UNC-104 PH HMW normalized to the UNC-104 PH LMW from the western blot in D. Results are shown as Mean ± SD (N=3 biological repeats represented as filled circles). *p<0.05 (Unpaired student’s t-test two-tailed). E) Western blot analyses from anti-myc IP enriched fractions of worms expressing fragments of various deletions to the N terminus of the PH domain marked with Δ1(Δ36 aa), Δ2(Δ20 aa), Δ3(Δ14 aa), Δ4(Δ20 aa). The input served as a loading control probed with β-Actin. Molecular weights (in kDa) are marked with arrows.

To find the site of this modification near the PH domain, we created four sub-deletions of the PH domain-containing fragment 4 (Δ1, Δ2, Δ3, Δ4) all of which contain lysine residues [Fig. 4E, Table S7]. The presence of the ~8kDa higher molecular weight band persisted upon deletion of regions Δ2, Δ3, Δ4 [Fig. 4E, Table S7]. However, upon deletion of the first 36 aa (Δ1) of fragment 4 [Fig. S3D], the higher molecular weight band is not seen [Fig. 4E, Table S7]. Thus, fragment 4 is likely modified in the Δ1 region by a mono-ubiquitin/ubiquitin-like modifier, leading to the presence of the higher molecular weight band.

To summarize, *fbxb-65* likely regulates the modification of fragment 4 region of the UNC-104 motor leading to the presence of a higher molecular weight band. The modification likely occurs in the 1386-1421 aa region, N-terminal to the cargo binding PH domain. FBXB-65 may either directly modify UNC-104 or regulate UNC-104’s modification through other E3 ligases (Liu and Xirodimas, 2010; Ohh et al., 2002; Ohki et al., 2009; Oved et al., 2006; Tatham et al., 2008; Uzunova et al., 2007).

### VI. *fbxb-65* RNAi regulates RAB-3 transport to neuronal distal ends

UNC-104 is essential for transporting heterogeneous pre-SVs out of the neuronal cell body (Hall and Hedgecock, 1991). Since we observe an increase in UNC-104 anterograde flux and particle intensity, we examined if knockdown of *fbxb-65* leads to increased anterograde transport of its pre-SV cargo, RAB-3.

Compared to control animals, *fbxb-65* RNAi animals show an increased number of GFP::RAB-3 puncta at the PLM distal end [Fig. 5A]. GFP::RAB-3 labelled cargo redistribute towards the end of the neuronal process in *fbxb-65* RNAi animals (~8 puncta per 40 μm) with fewer puncta in the proximal process (~5 puncta per 40 μm) compared to control animals (~6 puncta per 40 μm and ~7.5 puncta per 40 μm respectively) [Fig. 5B,D, Table S8]. In contrast to the neuronal process, there was no significant change in GFP::RAB-3 intensity at the synapses between *fbxb-65* RNAi and the control [Fig. 5C, Table S8]. Therefore, *fbxb-65* RNAi leads to the accumulation of UNC-104 cargo GFP::RAB-3 towards the distal ends without affecting its distribution at the synapses of the PLM neuron.

**Figure 5.**
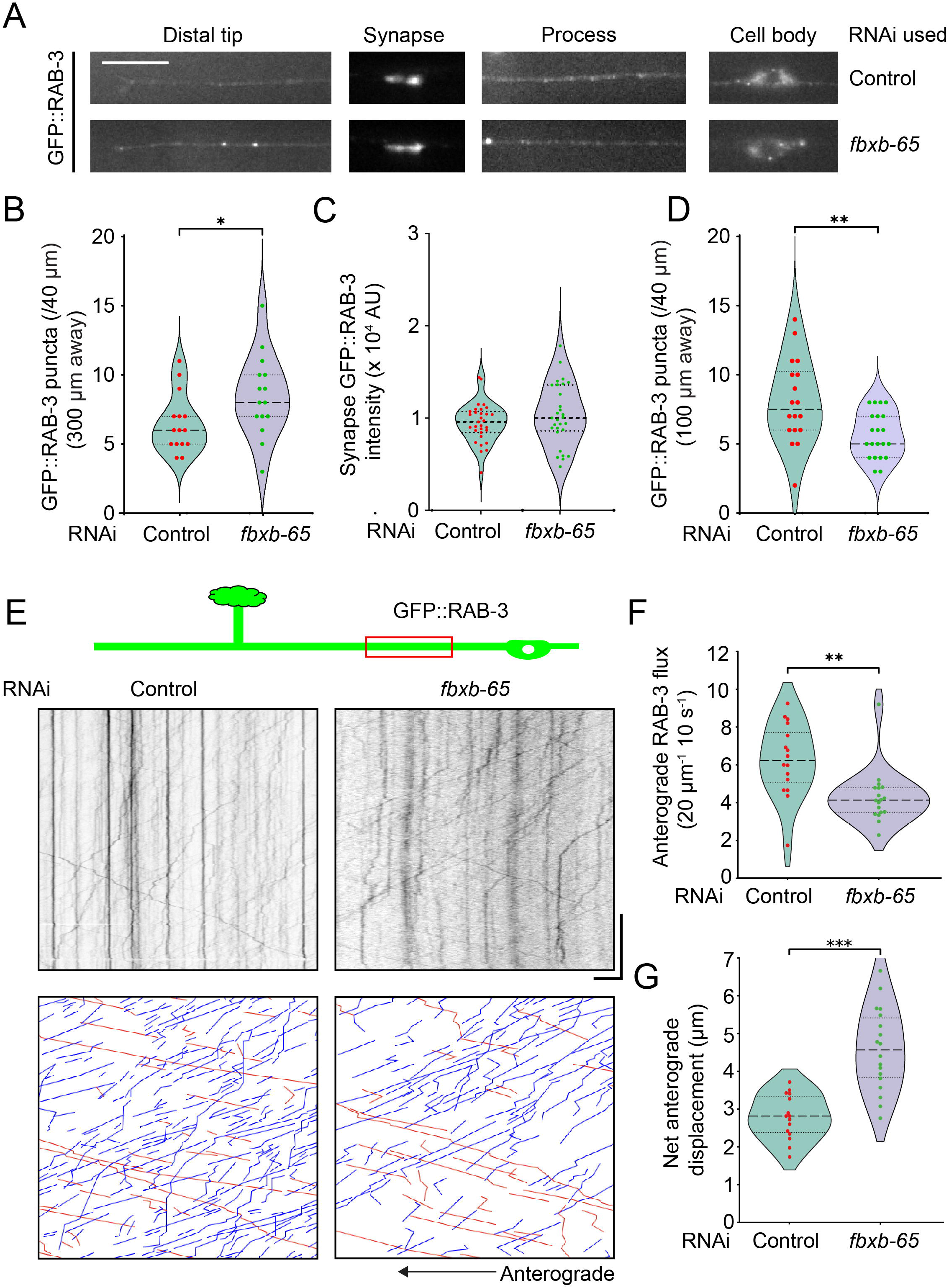
*fbxb-65* RNAi results in increased transport of RAB-3 cargo towards distal regions of the neuronal process. A) Representative images showing GFP::RAB-3 intensity at various regions in the PLM neuron including the cell body, proximal process (70 µm from the cell body), synapses, and the distal end in the strain TT2775 treated with either control or *fbxb-65* RNAi. Scale bar 10 µm. B) Number of GFP::RAB-3 puncta observed in a 40 µm region 300 µm away from the cell body in the strain TT2775 treated with either control (N=15) or *fbxb-65* (N=15) RNAi. Data are represented as violin plots with individual data points along with the median (dashed line), 25^th^ and 75^th^ percentile marked (dotted lines). *p<0.05 (Unpaired Student’s *t*-test two-tailed). C) Intensity of GFP::RAB-3 at the synapse assessed from the strain TT2775 treated with either control (N=15, 2 synapses each) or *fbxb-65* (N=15, 2 synapses each) RNAi. Data are represented as violin plots with individual data points along with the median (dashed line), 25^th^ and 75^th^ percentile marked (dotted lines); ns, non-significant (Unpaired Student’s *t*-test two-tailed). D) Number of GFP::RAB-3 puncta observed in a 40 µm region 100 µm away from the cell body in the strain TT2775 treated with either control (N=15) or *fbxb-65* (N=15) RNAi. Data are represented as violin plots with individual data points along with the median (dashed line), 25^th^ and 75^th^ percentile marked (dotted lines). **p<0.01 (Unpaired Student’s *t*-test two-tailed). E) Representative kymographs for GFP::RAB-3 in a region 70 µm from the cell body in the strain TT2775 treated with either control or *fbxb-65* RNAi along with their respective manually-traced annotations shown at the bottom. Blue lines represent a net anterograde motion, while red lines represent a net retrograde motion. Scale x-axis 10 µm, y-axis 30 s. F) Number of anterograde GFP::RAB-3 trajectories normalized to a 20 µm and 10 s region of the kymograph assessed from the strain TT2775 treated with either control (N=16) or *fbxb-65* (N=18) RNAi. Data are represented as violin plots with individual data points along with the median (dashed line), 25^th^ and 75^th^ percentile marked (dotted lines). **p<0.01 (Unpaired Student’s *t*-test two-tailed). G) Net displacement of anterogradely moving GFP::RAB-3 trajectories assessed from the strain TT2775 treated with either control (N=16) or *fbxb-65* (N=18) RNAi. Data are represented as violin plots with individual data points along with the median (dashed line), 25^th^ and 75^th^ percentile marked (dotted lines). ***p<0.001 (Unpaired Student’s *t*-test two-tailed).

Previous studies *in vitro* suggest higher numbers and increased activity of motors on cargo can be associated with longer single runs of cargo (Furuta et al., 2013; Vershinin et al., 2007). Thus, we examined movement properties of UNC-104 dependent GFP::RAB-3 tagged pre-SVs in the PLM [Fig. 5E]. There is an increase in the average anterograde run length and the fraction of GFP::RAB-3 with run lengths >2 μm in *fbxb-65* RNAi compared to the control [Figs. 5G, S4B, Tables S8, S15]. GFP::RAB-3-tagged vesicles have a similar velocity (0.6 μm s^-1^) and reduced flux (~6 vesicles per 20 μm per 10 s) compared to that in control animals [Figs. 5F, S4A, Tables S8, S15].

To summarize, in *fbxb-65* RNAi, we observe a decrease in GFP::RAB-3 puncta closer to the cell body and an increase in puncta farther from the cell body. GFP::RAB-3-carrying pre-SVs shows increased run-length in *fbxb-65* RNAi; that may arise from increased numbers of motors on the cargo surface. It is likely that the increased number of moving UNC-104::GFP puncta is associated with GFP::RAB-3 labelled synaptic vesicle cargo, influencing both their movement and distribution in the neuron.

### VII. Null mutants of *fbxb-65* and *unc-104(Δ1)* phenocopy *fbxb-65* knockdown

To verify the *fbxb-65* RNAi phenotypes, we generated a line containing CRISPR-mediated deletion of *fbxb-65*, named *fbxb-65(syb7320),* and hereon referred to as *fbxb-65(0)*. We also generated a CRISPR-mediated deletion line wherein the 36 aa (Δ1) near the UNC-104 PH domain (see above) that is required for FBXB-65 dependent modification is deleted. These mutants are named *unc-104(syb7293)*, hereon referred to as *unc-104(Δ1)*. We assessed the effects of these mutants on UNC-104::GFP CRISPR knock-in to examine if the endogenous motor was affected similarly to that observed in the TRN specific transgenically expressed UNC-104::GFP.

The *fbxb-65(0)* animals show accumulation of CRISPR knock-in UNC-104::GFP at the PLM distal end, like that observed with *fbxb-65* RNAi [Fig. S5A]. In addition, UNC-104::GFP knock-in accumulates at distal ends of the ALM neuron and other neurons that terminate in the head [Fig. 6A, Table S9]. Similar to *fbxb-65* RNAi, there is an increase in the average number (~1.2 per 20 μm per 10 s) and particle intensity (11 AU) of moving UNC-104::GFP knock-in puncta in *fbxb-65(0)* compared to that seen in UNC-104::GFP knock-in animals (~0.9 per 20 μm per 10 s and 7 AU, respectively) [Fig. 6C,D, Table S9]. The run lengths of UNC-104::GFP knock-in trajectories were largely unaltered (~12.5 μm) as seen in *fbxb-65* RNAi [Fig. S5B, Table S16]. Similar to *fbxb-65* RNAi, the intensity histogram of UNC-104::GFP knock-in trajectories in *fbxb-65(0)* animals are right shifted compared to UNC-104::GFP knock-in animals with *μ_control_* > *μ_fbxb-65_* [Fig. S5C,D]. Thus, UNC-104::GFP knock-in with *fbxb-65(0)* phenocopies the intensity distribution and movement properties of UNC-104::GFP observed in *fbxb-65* RNAi.

**Figure 6.**
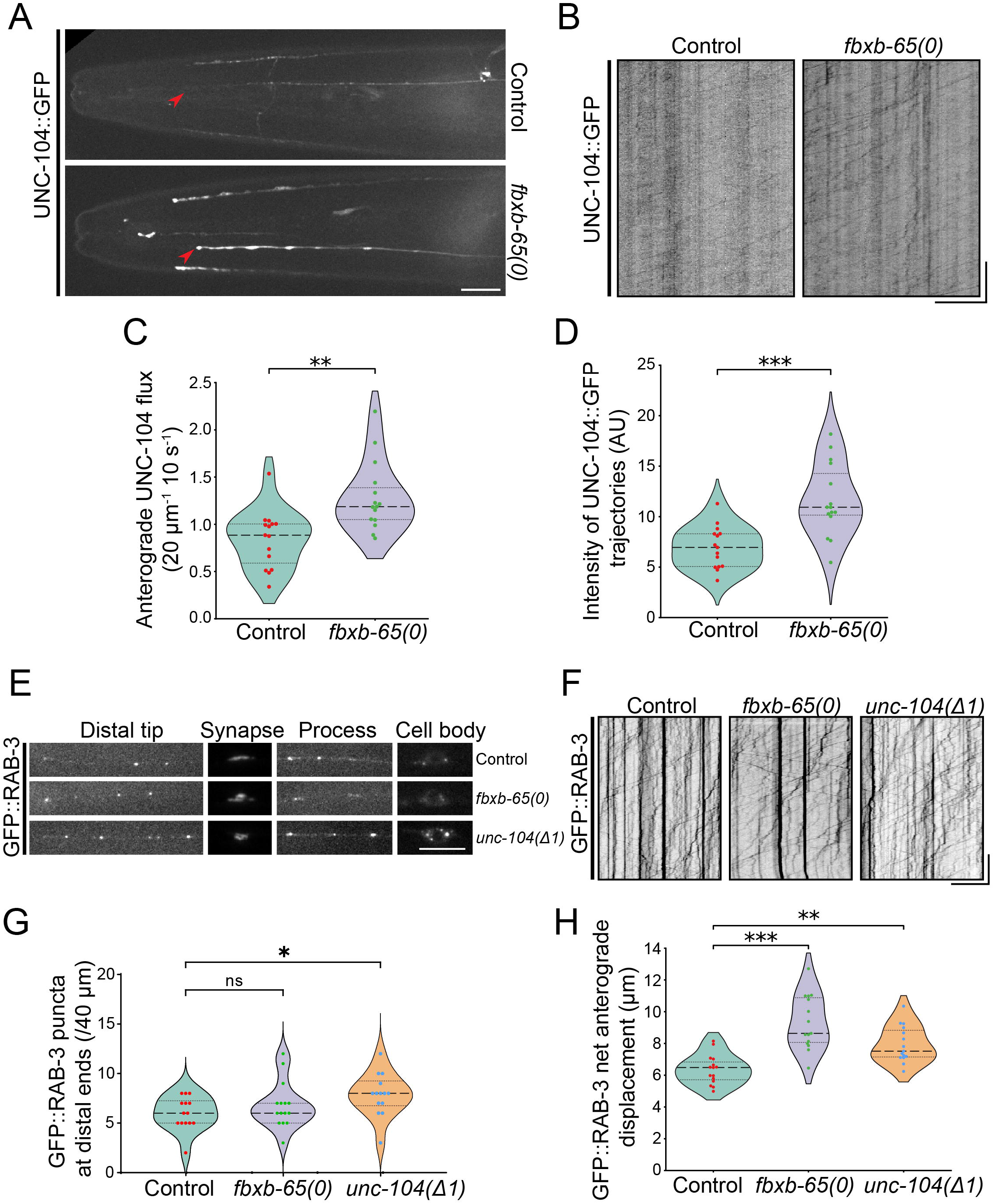
*fbxb-65(0)* and *unc-104(Δ1)* show similar phenotypes as *fbxb-65* RNAi. A) Representative images of the head of *C. elegans* showing the ALM tip (marked with a red arrow red) along with other neuronal ends in the vicinity in either the control *xdKi3* (endogenously-tagged UNC-104::GFP) or *fbxb-65(0)*. Scale 10 µm. B) Representative kymographs showing *xdKi3* at 70 µm from the cell body either in control or with *fbxb-65(0)* Scale x-axis 10 µm, y-axis 20 s. C) Number of anterograde UNC-104::GFP trajectories normalized to a 20 µm and 10 s region of the kymograph in either *xdKi3* control (N=15) or with *fbxb-65(0)* (N=15). Data are represented as violin plots with individual data points along with the median (dashed line), 25^th^ and 75^th^ percentile marked (dotted lines). ***p<0.001 (Unpaired Student’s *t-*test two-tailed). D) Background subtracted intensity of anterograde UNC-104::GFP trajectories assessed in either *xdKi3* control (N=15) or with *fbxb-65(0)* (N=15). Data represented as violin plots with individual data points along with the median (dashed line), 25^th^ and 75^th^ percentile marked (dotted lines). ***p<0.001 (Unpaired Student’s *t*-test two-tailed). E) Representative images of the cell body, proximal process 100 µm from the cell body, synapse or distal end of the PLM expressing GFP::RAB-3 in either control, *fbxb-65(0)* or *unc-104(Δ1)* animals. Scale 10 µm. F) Representative kymographs of GFP::RAB-3 in a region 70 µm from the cell body in either control, *fbxb-65(0)* or *unc-104(Δ1)* animals. Scale x-axis 10 µm, y-axis 20 s. G) Number of GFP::RAB-3 puncta observed in a 40 µm region 300 µm away from the cell body in either control (N=14), *fbxb-65(0)* (N=15) or *unc-104(Δ1)* (N=14) animals. Data are represented as violin plots with individual data points along with the median (dashed line), 25^th^ and 75^th^ percentile marked (dotted lines). *p<0.05 (one-way ANOVA with Dunnett’s multiple comparisons test). H) Net displacement of anterogradely moving GFP::RAB-3 trajectories assessed in either control (N=15), *fbxb-65(0)* (N=15) or *unc-104(Δ1)* (N=15) animals. Data are represented as violin plots with individual data points along with the median (dashed line), 25^th^ and 75^th^ percentile marked (dotted lines). **p<0.01 ***p<0.001 (one-way ANOVA with Dunnett’s multiple comparisons test).

We also assessed if *fbxb-65(0)* led to a change in GFP::RAB-3 transport. The average run length of GFP::RAB-3 in *fbxb-65(0)* (~8.6 μm) is increased compared to that in wild type animals (~6.4 μm) [Fig. 6H, Table S9]. GFP::RAB-3 run lengths in both *fbxb-65(0)* and *fbxb-65* RNAi is increased compared to their respective controls [Figs. 5G, 6H, Tables S8, S9]. The flux (~2.2 per 20 μm per 10 s) and velocity (~0.9 μm s^-1^) of GFP::RAB-3 in *fbxb-65(0)* were largely similar to those seen in wild type animals [Fig. S5E,F, Table S16].

We assessed whether the deletion of 36 aa near the PH domain of UNC-104 that is the region modified by FBXB-65, also alters GFP::RAB-3 distribution and transport properties akin to those seen in *fbxb-65(0).* The number of GFP::RAB-3 puncta at the distal ends of the neuron (~8 per 40 μm) and GFP::RAB-3 run length (~7.5 μm) is increased in *unc-104(Δ1)* compared to that seen in wild type animals (~6 per 40 μm and ~6.4 μm, respectively) [Fig. 6G,H, Table S9]. The velocity and flux of GFP::RAB-3 are largely unchanged in *unc-104(Δ1)* compared to those in wild type animals, similar to what we observe in *fbxb-65(0)* animals [Fig. S5E,F, Table S16]. Thus, both *fbxb-65(0)* and *unc-104(Δ1)* have similar effects on GFP::RAB-3-associated pre-SV movement, similar to that seen in *fbxb-65* RNAi.

To summarize, the *fbxb-65(0)* mutant phenocopies *fbxb-65* RNAi, with distal end accumulation of UNC-104 and increased intensity of moving UNC-104::GFP trajectories. Both *fbxb-65(0)* and *unc-104(Δ1)* show increased GFP::RAB-3-associated cargo run length, with no significant change in velocity or flux of moving GFP::RAB-3 particles, similar to GFP::RAB-3 movement observed in *fbxb-65* RNAi animals. Furthermore, the number of GFP::RAB-3-associated cargo at the neuronal ends increases in *unc-104(Δ1)*. These data together suggest that loss of UNC-104 modification owing to loss of *fbxb-65* may permit increased UNC-104 association with cargo, thereby increasing cargo run length and redistribution of both motors and cargo to the end of the neuron.

### VIII. *fbxb-65* can increase the extent of axonal localization of synaptic vesicle proteins in pre-SV transport-defective mutants

The distance cargo travels in the axon depends on net motor activity (Hayashi et al., 2019; Niwa et al., 2017). In partial loss of function *unc-104(e1265tb120)* mutants with lower levels of motors, pre-SV associated proteins travel into the axon but do not reach the synapse (Kumar et al., 2010). In *sam-4* mutants with both lower UNC-104 levels and accumulation of the motor in the cell body, pre-SV associated proteins accumulate near the cell body and very little reaches the synapse (Zheng et al., 2014). UNC-104 overexpression can suppress the pre-SV transport defects of mutants of the UNC-104 regulator *sam-4* (Zheng et al., 2014). We tested if *fbxb-65*, which likely controls the number of UNC-104 on pre-SVs, phenocopies UNC-104 overexpression and rescues both *unc-104(e1265tb120)* and *sam-4(js415)* phenotypes.

In *unc-104(e1265tb120)*, GFP::RAB-3 is present up to ~60 μm into the PLM neuronal process unlike in wild type animals, where GFP::RAB-3 is present along the entire ~350 μm length of the PLM neuronal process [Figs. 5A, 7A]. In the double mutant *unc-104(e1265tb120)*; *fbxb-65(0)* animals, GFP::RAB-3 travels further into the neuronal process and is present until ~80 μm [Fig. 7A]. Likewise, in *sam-4(js415)* treated with control RNAi, GFP::RAB-3 accumulates close to the cell body and only ~40% animals show GFP::RAB-3 at the branch point (200–300 μm from the cell body) or at synapses [Fig. 7C] (Zheng et al., 2014). Overexpression of UNC-104 in *sam-4(js415)* reduces GFP::RAB-3 accumulation near the cell body [Fig. 7D, Table S10] and leads to an increased number of animals with GFP::RAB-3 at the branch point (~80%) compared to *sam-4(js415)* [Fig. 7E, Table S10] (Zheng et al., 2014). In *sam-4(js415)* treated with *fbxb-65* RNAi, we see a similar decrease in GFP::RAB-3 accumulation near the cell body and an increase in number of animals with GFP::RAB-3 present at the branch point (~87%) [Fig. 7D,E, Table S10]. Likewise, in *sam-4(js415)*; *fbxb-65(0)* double mutant animals, the accumulation of GFP::RAB-3 puncta in the proximal neuronal process is reduced compared to that seen in *sam-4(js415)* animals [Fig. 7F,G, Table S10]. Thus, consistent with increased UNC-104 on the cargo surface, loss or reduction of *fbxb-65* leads to presence of GFP::RAB-3 further into the neuronal process in the *sam-4(js415)* background.

**Figure 7.**
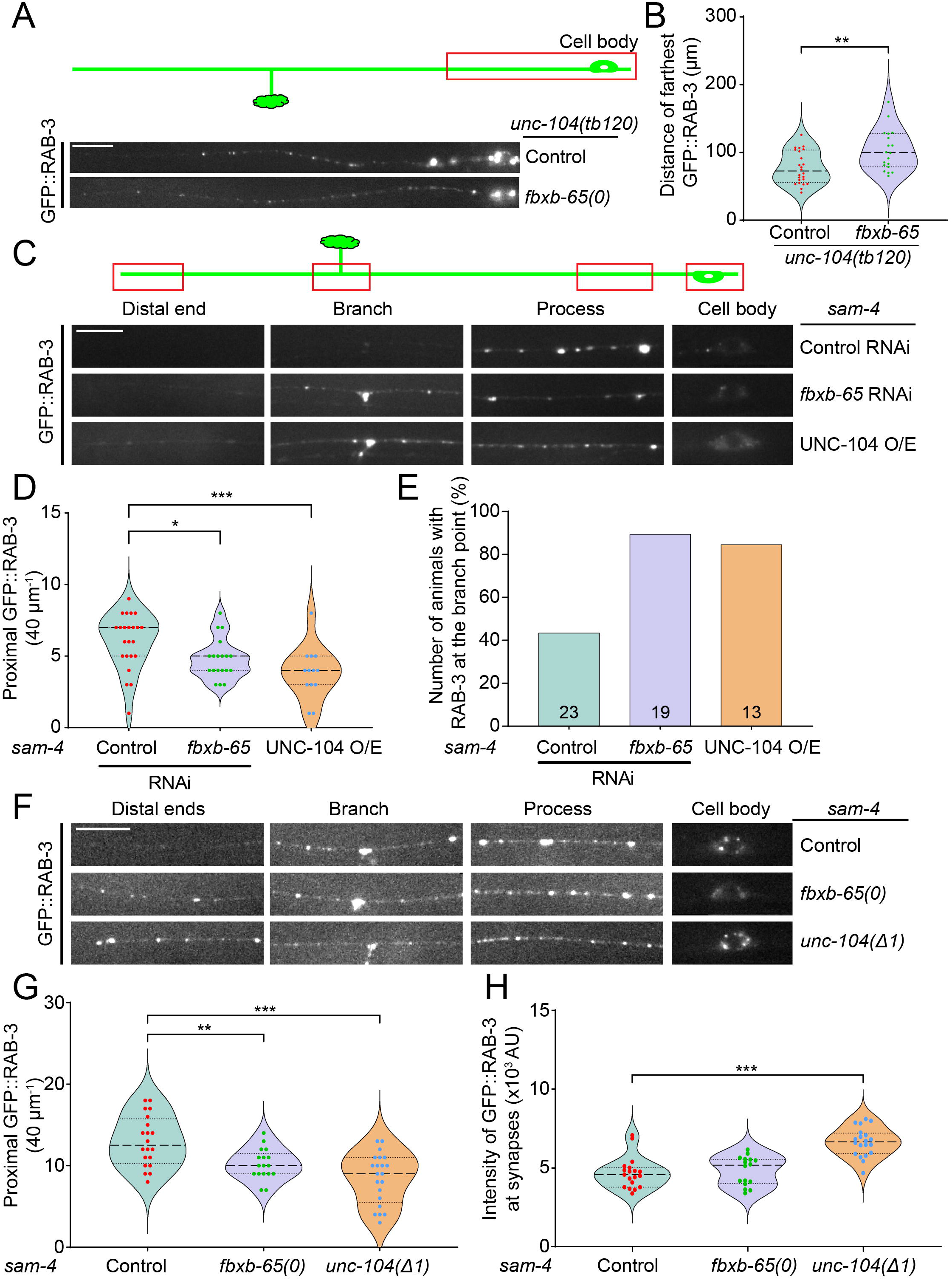
*fbxb-65* regulates extent of GFP::RAB-3 in *unc-104(e1265tb120)* and *sam-4(js415)* A) Schematic showing region of imaging along with representative images of the first 100 µm expressing GFP::RAB-3 with either the single mutant *unc-104(e1265tb120)* or the double mutant *unc-104(e1265tb120)*; *fbxb-65(0)*. Scale 10 µm. B) Farthest distance of GFP::RAB-3 puncta observed in the single mutant *unc-104(e1265tb120)* (N=24) or the double mutant *unc-104(e1265tb120)*; *fbxb-65(0)* (N=17). Data are represented as violin plots with individual data points along with the median (dashed line), 25^th^ and 75^th^ percentile marked (dotted lines). **p<0.01 (Unpaired Student’s *t*-test two-tailed). C) Schematic showing regions of imaging including the cell body, proximal process (70 µm from the cell body), synapses, and the distal end along with their representative images of GFP::RAB-3 in the strain TT3186 treated with either control or *fbxb-65* RNAi or the strain TT3230 that has a pan-neuronal overexpression of UNC-104. Scale bar 10 µm. D) Number of GFP::RAB-3 puncta observed in a 40 µm region 100 µm away from the cell body in the strain TT3186 treated with either control (N=23) or *fbxb-65* (N=19) RNAi or the strain TT3230 (N=13). Data are represented as violin plots with individual data points along with the median (dashed line), 25^th^ and 75^th^ percentile marked (dotted lines). *p<0.05 **p<0.01 (one-way ANOVA with Dunnett’s multiple comparisons test). E) Number of animals represented as a percentage that have GFP::RAB-3 puncta visible at the distal end of the neuronal process in the strain TT3186 treated with either control (N=23) or *fbxb-65* (N=19) RNAi or the strain TT3230 (N=13). F) Representative images of animals expressing GFP::RAB-3 with either the single mutant *sam-4(js415)*, the double mutant *sam-4(js415)*; *fbxb-65(0)* or the double mutant *sam-4(js415) unc-104(Δ1)*. Scale bar 10 µm. G) Number of GFP::RAB-3 puncta observed in a 40 µm region 100 µm away from the cell body in either the single mutant *sam-4(js415)* (N=20), the double mutant *sam-4(js415)*; *fbxb-65(0)* (N=17) or the double mutant *sam-4(js415) unc-104(Δ1)* (N=21). Data are represented as violin plots with individual data points along with the median (dashed line), 25^th^ and 75^th^ percentile marked (dotted lines). **p<0.01 ***p<0.001 (one-way ANOVA with Dunnett’s multiple comparisons test). H) Intensity of GFP::RAB-3 at synapses measured in either the single mutant *sam-4(js415)* (N=20), the double mutant *sam-4(js415)*; *fbxb-65(0)* (N=17) or the double mutant *sam-4(js415) unc-104(Δ1)* (N=21). Data are represented as violin plots with individual data points along with the median (dashed line), 25^th^ and 75^th^ percentile marked (dotted lines). ***p<0.001 (one-way ANOVA with Dunnett’s multiple comparisons test).

We used *unc-104(Δ1)* as an allele of *unc-104* that is independent of *fbxb-65*, potentially encoding UNC-104 motors that associate in greater numbers with cargo. In the double mutant *unc-104(Δ1) sam-4(js415)*, there are fewer GFP::RAB-3 puncta in the proximal process [Fig. 7F,G, Table S10] with an increase in synaptic GFP::RAB-3 [Figs. 7H, S5G, Tables S10, S16] compared to *sam-4(js415)* animals. Thus, the motor encoded by *unc-104(Δ1)* may associate more with pre-SVs since they are incapable of being modified by FBXB-65. The FBXB-65 independent UNC-104 suppresses *sam-4(js415)* and leads to increased GFP::RAB-3 at synapses.

To summarize, increasing the number of motors capable of associating with pre-SVs either by removing/reducing FBXB-65 or making UNC-104 independent of FBXB-65 both lead to improved pre-SV transport in *sam-4(js415)* and *unc-104(e1265tb120)* animals. The modification of UNC-104 by FBXB-65 may functionally titrate the levels of motors on the pre-SV cargo surface and regulate pre-SV cargo distribution.

### IX. *fbxb-65*-regulated UNC-104 modification facilitates turning of synaptic vesicles into the synaptic branch

Levels and types of motors have been shown to be critical in navigating complex microtubule arrangements (Bergman et al., 2018; Tymanskyj et al., 2022). The PLM TRN has a synaptic branch that drops from the main process [Fig. 8A]. pre-SVs turning into the synaptic branch depends on numbers of UNC-104 motors (Vasudevan et al., 2023 preprint). UNC-104 overexpression leads to an increased incidence of pre-SVs going straight to the non-synaptic neuronal terminus in the PLM neuron (Vasudevan et al., 2023 preprint). Since the numbers of UNC-104 on the per-SV cargo surface are likely increased in *fbxb-65(0)* and *unc-104(Δ1)*, we tested if they controlled the behavior of vesicles at PLM TRN branch points.

**Figure 8.**
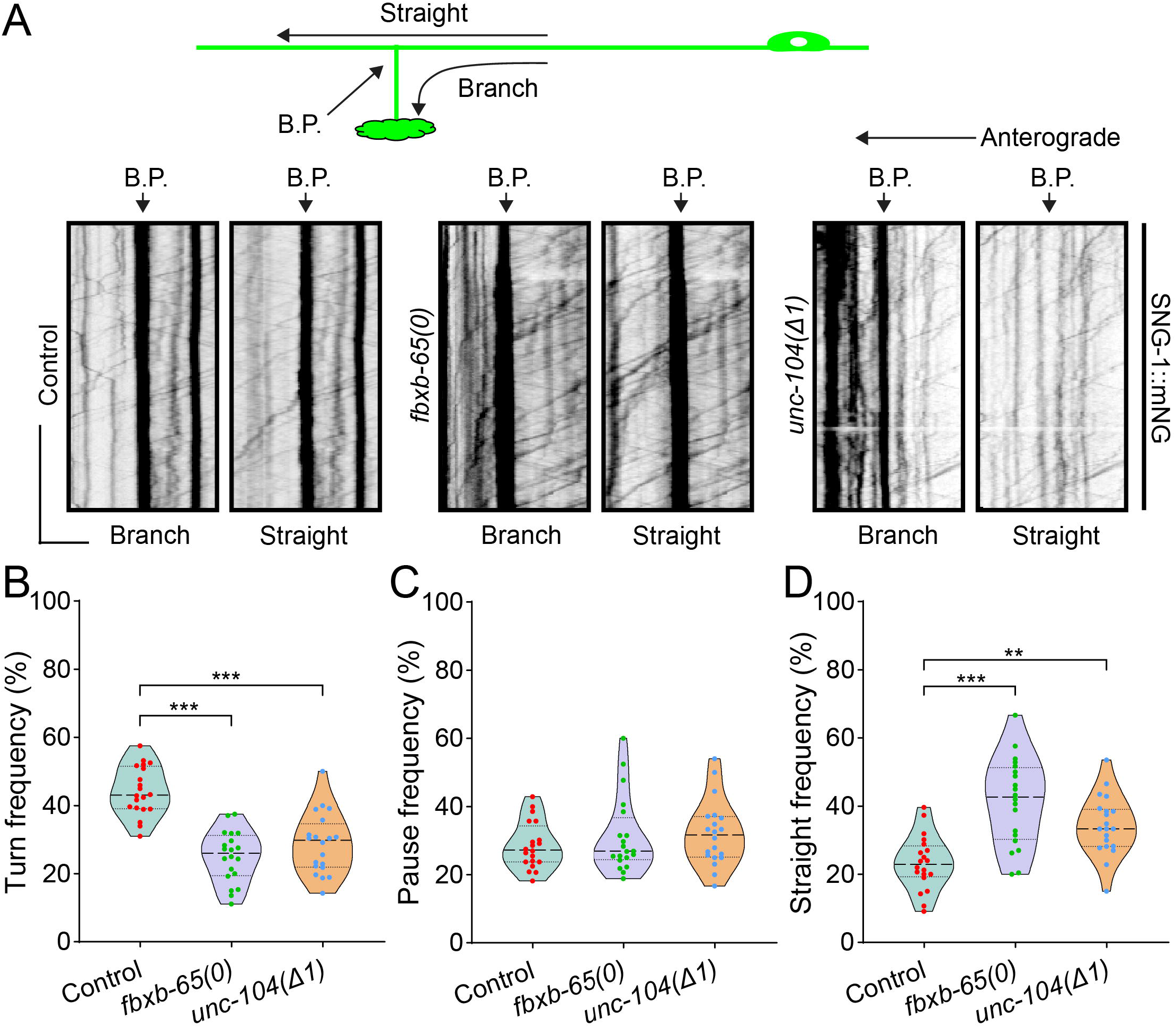
*fbxb-65* RNAi increases flux and intensity of UNC-104 single particle tracks compared to control. A) Representative kymographs for SNG-1::mNeonGreen traced either along the branch or along the straight process in either control (N=20), *fbxb-65(0)* (N=20), or *unc-104(Δ1)* (N=20) as shown in the schematic. The branch point (B.P.) marked by arrows in the kymograph and the schematic. Scale x-axis 5 µm, y-axis 30 s. B) Percentage of SNG-1::mNeonGreen vesicles turning into the branch out of all vesicles that approach the branch point anterogradely in either control (N=20), *fbxb-65(0)* (N=20), or *unc-104(Δ1)* (N=20). Data are represented as violin plots with individual data points along with the median (dashed line), 25^th^ and 75^th^ percentile marked (dotted lines); ***p<0.001 (one-way ANOVA with Dunnett’s multiple comparisons test). C) Percentage of SNG-1::mNeonGreen vesicles pausing at the branch out of all vesicles that approach the branch point anterogradely in either control (N=20), *fbxb-65(0)* (N=20), or *unc-104(Δ1)* (N=20). Data are represented as violin plots with individual data points along with the median (dashed line), 25^th^ and 75^th^ percentile marked (dotted lines). D) Percentage of SNG-1::mNeonGreen vesicles going straight to the non-synaptic distal end out of all vesicles that approach the branch point anterogradely in either control (N=20), *fbxb-65(0)* (N=20), or *unc-104(Δ1)* (N=20). Data are represented as violin plots with individual data points along with the median (dashed line), 25^th^ and 75^th^ percentile marked (dotted lines); **p<0.01 ***p<0.001 (one-way ANOVA with Dunnett’s multiple comparisons test).

In *fbxb-65(0)*, there is an increased number of SNG-1::mNeonGreen marked pre-SVs going straight toward the non-synaptic distal end along with a concomitant decrease in vesicles turning toward the synapse compared to those in wild type animals [Fig. 8B,D, Table S11]; *unc-104(Δ1)* also shows similar phenotypes [Fig. 8B,D, Table S11]. The percentage of vesicles pausing at the branch point are similar across wild type, *fbxb-65* and *unc-104(Δ1)* animals [Fig. 8C]. These phenotypes resemble UNC-104 overexpression in the PLM (Vasudevan et al., 2023 preprint). Therefore, loss in *fbxb-65*-mediated regulation of UNC-104 modification may mistarget synaptic vesicles across branch points away from their destination, the synapse.

## Discussion

Post-translational modification of motors has been shown to regulate motor activity (Liang et al., 2014), motor–cargo interactions (Guillaud et al., 2008; Matthies et al., 1993), and motor conformation (Espeut et al., 2008). Here, we show that a modification near the cargo-binding PH domain of UNC-104 is regulated by a putative E3 ligase FBXB-65, and that this modification potentially reduces the number of motors on the cargo surface thereby altering anterograde motor and cargo movement. The significance of FBXB-65-mediated regulation becomes apparent in situations where cargo transport is compromised. Our findings suggest that a post-translational modification can regulate motor–cargo binding to regulate the number of motors on the cargo surface. Additionally, our theoretical model suggests that cooperative binding of UNC-104 may account for some of our observations.

An increased number of kinesins on cargo have been shown to increase cargo run lengths *in vitro,* although not *in vivo* (Beeg et al., 2008; Shubeita et al., 2008; Wilson et al., 2021). *In vivo* increases in cargo run lengths and flux have been shown to arise from post-translational modifications, such as phosphorylation, of either the motor or its adapter (Padzik et al., 2016; Prowse et al., 2023). The UNC-104 motor binds to its cargo through phosphatidylinositol 4,5-bisphosphate (PIP_2_) (Klopfenstein and Vale, 2004; Kumar et al., 2010), and on average, 132 phosphatidylinositol molecules have been reported to be present on each synaptic vesicle (Takamori et al., 2006). *Dictyostelium* UNC-104 motors have been shown to cluster and display higher processivity with increasing levels of PIP_2_ (Klopfenstein et al., 2002). Building on these findings, using a theoretical model to explain the distribution of UNC-104 cluster intensity, we show that UNC-104 can bind cooperatively, resulting in higher motor intensities than predicted by a non-cooperative model. Consequently, the increased number of motors, likely on cargo, observed in *fbxb-65* animals may exhibit a greater propensity for clustering, thereby explaining the longer GFP::RAB-3 associated cargo run lengths [Fig. 5G]. Increased UNC-104 clustering may be mediated by proteins such as UNC-10, which acts as a linker between UNC-104 and RAB-3 associated pre-SVs (Bhan et al., 2020 preprint), SYD-2 and UNC-10, which are known to interact with and activate UNC-104 (Wagner et al., 2009; Wu et al., 2016), or by phospholipids, which may cluster on the cargo surface (Klopfenstein et al., 2002). The modification of UNC-104 by *fbxb-65* may regulate UNC-104 levels on the cargo surfagce by deterring UNC-104’s ability to bind to the cargo surface effectively.

Movement of synaptic vesicles can face challenges due to narrow and complex intracellular environments enroute to their destination, the synapse (Sabharwal and Koushika, 2019; Sood et al., 2018). Increased KIF1A/UNC-104 promotes synaptogenesis (Kondo et al., 2012) and improves behavior in aged animals (Li et al., 2016). The number of motors on the cargo surface is likely critical for both how far the cargo travels along the axon, how cargo navigate obstacles and steady-state distribution of cargo (Chen et al., 2020). Some KIF1A-associated neurological disorder afflicted patients carry gain-of-function KIF1A mutations that may lead to increased motor velocity or landing rates on microtubules, which lead to aberrant accumulation of synaptic vesicle proteins at neuronal distal ends (Chiba et al., 2019; Gabrych et al., 2019). Similarly, we observe increased GFP::RAB-3 puncta at distal ends and reduced GFP::RAB-3 puncta in the proximal neuronal process in *fbxb-65* RNAi [Fig. 5B,D]. The number of motors bound to cargo likely exists in a dynamic equilibrium with freely diffusing cargo-unbound motors in the vicinity, as suggested for Kinesin-1 (Blasius et al., 2013). Increased attachment of motor to cargo may lead to phenotypes similar to those seen in motors defective in auto-inhibition. Such regulation may, in part, be mediated by FBXB-65, which is present throughout the neuronal process [Fig. S1A]. In situations where cargo transport is compromised, for instance upon reduced cargo binding in UNC-104(D1497N M1540I) (Kumar et al., 2010; Maeder et al., 2014), or reduced cargo processivity in *sam-4(js415)* (Zheng et al., 2014), not having the modification can bypass these effects. However, unregulated UNC-104 pre-SV association upon reduced modification leads to mistargeting of pre-SV cargo away from synaptic regions [Fig. 8B,D]. Therefore, while numerous factors influence cargo run lengths and distribution *in vivo*, increased motors on the cargo surface may serve as a mechanism to overcome local obstacles, and regulation of motors on cargo may facilitate efficient transport of cargo to their destination.

Ubiquitin, initially discovered for its involvement in ATP-dependent protein degradation (Ciechanover et al., 1980; Hershko et al., 1980), plays a crucial role in regulating protein turnover in neurons (Speese et al., 2003). Previous studies have consistently observed an accelerated turnover of KIF1A/UNC-104, surpassing the average lifetime of other neuronal proteins and even other members of the kinesin family (Cohen et al., 2013; Fornasiero et al., 2018; Huang et al., 2020; Mathieson et al., 2018). A prior study demonstrates ubiquitin-dependent degradation of UNC-104 (Kumar et al., 2010). We propose that a ubiquitin or ubiquitin-like modifier serves as a regulator governing the association of UNC-104 with pre-SV cargo, thereby fine tuning the levels and activity of UNC-104 within neurons. Notably, these modifications, mediated by E3 ligases (Chu and Yang, 2011; Oved et al., 2006; Schmidt and Dikic, 2005), may elicit distinct outcomes in different neuronal compartments, such as the synapse versus the neuronal process, indicating potential cross-talk between various modification types.

In conclusion, we propose that UNC-104 levels on pre-SV surface are controlled by a FBXB-65 regulated modification near the UNC-104 PH domain. While increased UNC-104 levels on pre-SVs may bypass transport defective conditions, regulating UNC-104 levels is likely essential for correct targeting of pre-SVs to the synapse and preventing misaccumulation of pre-SVs towards the neuronal distal ends. The lack of appropriate targeting with too much transport may be detrimental to neuronal health (Gabrych et al., 2019).

## Materials and Methods

### Strain maintenance and generation

Worms were maintained on Nematode Growth Media (NGM) agar at 20 °C, and spotted with the *Escherichia coli* strain OP50 (Brenner, 1974). RNAi experiments were conducted by transferring adults to an NGM plate, supplemented with 100 μg μl^-1^ ampicillin and 1 mM IPTG (Biobasic Catalog IB0168), 1 day after seeding with dsRNA-expressing *E. coli* HT115 isolated from the Ahringer *C. elegans* RNAi feeding library or the control Empty Vector L4440 RNAi. The larval stage 4 (L4) hermaphrodite worms were used for all experiments unless stated otherwise. Transgenic worms were generated by microinjecting into N2 using an Eppendorf® FemtoJet microinjection system. Injection cocktails were composed of 50 ng μl^-1^ co-injection marker, the required concentration of reporter DNA, and made up to 200 ng μl^-1^ with pBluescript vector. The strains PHX7320 *fbxb-65(syb7320)* and PHX7293 *unc-104b(syb7293)* were generated by SunyBiotech’s CRISPR service. Other strains were created using crosses by standard approaches. All strains used are listed in Supplementary Table 1.

### Confocal imaging

L4 animals were taken and anesthetized with 5 mM tetramisole and laid on a 5% agar pad. Live imaging of UNC-104::GFP (TT2440) and GFP::RAB-3 (TT2775) RNAi strains was performed using an Olympus IX83 fitted with a Yokogawa CSU-X1 excited with a 488 nm solid-state laser at 15% of 1 mW laser power at objective using a 100×/1.4 N.A. DIC oil objective with a pixel size of around 129 nm per pixel and imaged with a Hamamatsu EM-CCD camera.

Live imaging of the strain *xdKi3* was done on an Olympus IX83 fitted with a Yokogawa CSU-W1 excited with a 473 nm solid-state laser at 15% of 3 mW laser power at objective using a 100×/1.4 N.A. DIC oil objective and imaged with a Prime BSI sCMOS camera configured with 2×2 binning leading to a pixel size of around 130 nm per pixel.

To generate kymographs, the movies were taken with an exposure time of 300 ms leading to a frame rate of 3 frames per second for a total time of around 3 min.

Static images of UNC-104::GFP and GFP::RAB-3 were taken on an epifluorescent IX73 scope illuminated with 100% output from a 120 W X-cite mercury-arc lamp and imaged using a Photometrics Evolve 512 EMCCD camera and a 100×/1.4 N.A. DIC oil objective with a pixel size of 158 nm per pixel and exposure of 150 ms.

### Generation of transgenic *C. elegans*

Transgenic lines were generated by following standard techniques (Stinchcomb et al., 1985) using an Olympus IX53 equipped with 20× and 40× air objectives, Narishige M-152 micromanipulator, and Eppendorf Femtojet II microinjector. The F2 progenies that inherited and stably expressed the extrachromosomal transgene with >70% transmission was used for biochemistry experiments. All strains generated are listed in Supplementary Table 1.

### Image analysis

All image panels used for representation and analysis of time lapse movies were generated using Fiji-ImageJ v1.52p (Schindelin et al., 2012). Experimental kymographs were built using KymoResliceWide v.0.5 plugin for ImageJ (https://github.com/ekatrukha/KymoResliceWide) using the average intensity measured across a polyline of width 3 in a region around 100 μm distal to the PLM cell body. The kymographs were then annotated to include all events with a meaningful velocity (<5 μm s^-1^ and >0.1 μm s^-1^) manually to derive estimates of flux and run lengths. A cargo is counted as moving if it has been displaced by at least 3 pixels in successive time frames.

For estimating the intensity of a single UNC-104::GFP trajectory, the average intensity of a trajectory was subtracted by the average intensity of the same trajectory three pixels above (corresponding to 1 second prior) on a kymograph with time progression vertically downwards. Regions of the trajectory that had a pause were neglected from the analysis.

### Stochastic model to explain the intensity of moving motor puncta

The evolution of the probability *P*(*n,t*) of a cargo bound to *n* motor proteins (MP) can be expressed in terms of the following Master equation (Gardiner, 1985)

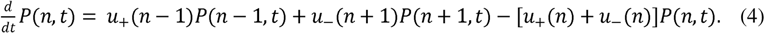

Here, *u*_+_(*n*) and *u*_−_(*n*) are the rates of binding and unbinding of an MP to the cargo, respectively. The rates can in general depend on the number of MPs already attached. The equation can be solved for steady-state using the boundary condition for *n* = 1 P attached state

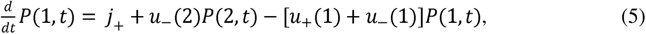

where *j*_+_ represents the diffusion-limited rate with which the first MP establishes cross-link between the cargo and the microtubule. At steady state, the flux of binding-unbinding of the first MP cross-linking the cargo to the microtubule must balance, i.e., *j*_+_ = *u*_−_(1)*P_S_*(1), with *P_S_*(1) denoting the steady-state probability. This relation, along with Eqn. 4 and Eqn. 5, lead to the pairwise balance condition

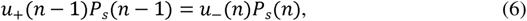

in a steady state. Here, *P_S_*(*n*) denotes the steady-state probability for cargo bound to *n* MPs. The exact steady-state solution can be obtained using a recurrence relation:

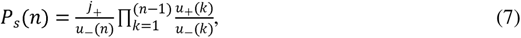

with the normalization condition 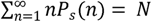, the total number of bound motors. Using the simplest assumption of a constant rate of unbinding *β*, one gets

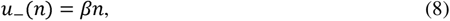

Here, we postulate that the binding rate is cooperative in nature, namely, the presence of MPs on cargo enhances the probability of more MPs attaching. Such a cooperative binding can be expressed up to linear order as:

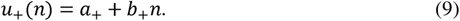

Using the expressions in Eqn. 8 and Eqn. 9 in Eqn. 6, and expanding up to linear order in 1/*N* and solving, we find a closed form solution for *P_S_*(*n*) given as (Chaudhuri et al., 2011):

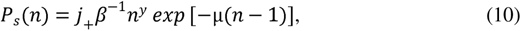

That obeys the boundary condition of cargo unbinding. Here *γ* = *a*_+_/*β* − 1, and *μ* = 1 − *b*_+_/*β*. The distribution in Eqn. 10 has a maximum at 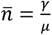, mean value 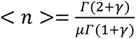 and variance 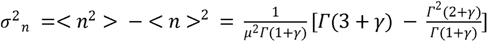, Experiments show that *γ* ≈ 1. Using *Γ* = 1 simplifies the above expressions to < *n* > = 2/*μ* and *σ*^2^*_n_* = 2/*μ*^2^.

We note that the form given in Eqn. 9 leads to a good agreement with the experimental distributions, both for control and *fbxb-65* RNAi. We further note that the best agreement is obtained for *γ* ≈ 1 for all cell types. This indicates that the bare binding rate *a*_+_/*β* ≈ 2 in units of *β*. We also note that the parameter *μ* > 0, which indicates that the coefficient of cooperative binding rate 0 < *b*_+_/*β* < 1. Note that in the absence of cooperative binding (*b*_+_ = 0), the exponential decay constant *μ* = 1 will be independent of the cell type. In contrast, experiments clearly show a strong cell-type dependence in the decay constant. This further shows that our hypothesis of cooperative binding is crucial in describing the distribution of bound MPs. Upon comparing the *μ* values obtained by fitting the experimental distributions with Eqn. 9, we note that *μ_control_* > *μ_fbxb-65_*. This indicates:

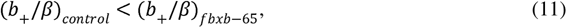

i.e., the cooperative binding increases for *fbxb-65* RNAi compared to the control. Further, the most parsimonious explanation that fits a change in µ is that *b*_+_ alters but not β, since *γ* ≈ 1 (*a*_+_. *β* ≈ 2) fits most of our data. Therefore, *fbxb-65-*mediated modification likely regulates recruitment of UNC-104 to cargo (altered *b*_+_) without altering UNC-104’s detachment kinetics (similar β).

### Ablation assay

The imaging was performed using an LSM 710 system with TT2440 worms anesthetized in 5 mM tetramisole on a 5% agar with a 63×/1.4 DIC M27 Oil objective at 3× zoom with a pixel size of 90 nm with a 488 nm Argon laser. Images were acquired every 500 ms in the ALM and PLM around 70 μm or 280 μm away from the cell body. Ablation was done using the Zen software and a femtosecond MaiTai Ti: Sapphire laser (Spectra-Physics) mode-locked at 800 nm at 80% power (400 mW maximum at objective) with 20 iterations post acquiring 5 pre-bleach images. The imaging was done for at least 200 frames post-bleach.

Post-acquisition, movies were analyzed using FIJI, by drawing a rectangular ROI of approximately 1×2 μm to cover the proximal and distal cut sites. The intensity at each time point was exported along with the ROI of bleaching and the ablation time point. The data were then normalized to pre-ablation intensity values and plotted with an in-house script generated in Python.

### qPCR estimation of RNA levels

RNA was isolated from TT3185 worms using TRIzol reagent (Thermo Fisher Scientific, catalog 15596018). Briefly, the worm pellet was freeze-cracked and RNA was extracted using a standard protocol (Rio et al., 2010). The isolated RNA (1 μg) was converted to cDNA using Superscript IV RT polymerase (Thermo catalog 18090010) according to the manufacturer’s protocol using random hexamers. The resultant cDNA was diluted 1:20 and used for qPCR reactions.

For qPCR reactions, primer efficiency was first calculated by amplifying cDNA template with KAPA 2× SYBR Master mix (Roche catalog 07959362001) and quantifying the SYBR intensity using a Roche LightCycler 480 at three different concentrations of cDNA 10-fold apart (1/2, 1/20, 1/200 dilutions). Only primers with an efficiency greater than 95% were used subsequently. For each qPCR reaction, controls for N2 and actin were set as negative control and standard, respectively. Each reaction was set in triplicates. The final fold change was determined using the ΔΔCt method. All primers are listed in Supplementary Table 2.

### Cloning of UNC-104 fragments

UNC-104 fragments were divided into four different fragments by performing PCR of the pSN8 construct (Niwa et al., 2016). Primers (Supplementary Table 2) were generated to clone a 5× MYC tag in the C-terminal of the fragments using in-fusion PCR. The combined fragment with MYC tag was then ligated with the vector backbone containing a pan-neuronal *rab-3* promoter pHW393 (*Prab-3*::GAL4-SK(DBD)::VP64::*let-858* 3’UTR), which was a gift from Paul Sternberg (Addgene plasmid # 85583; http://n2t.net/addgene:85583; RRID: Addgene_85583) (Wang et al., 2017). Further deletions in the PH domain-containing fragments were made using PCR followed by *in vivo* recombination in the *E. coli* strain DH5α. All reagents available upon request. All plasmids along with their construction details are listed in Supplementary Table 3.

### Biochemistry for motor ubiquitination

Worms were harvested from three almost-starved 60 mm NGM dishes by washing thrice in M9 buffer. Protein preparation from a mixed population of worms was adapted from a previous study (Shaham, 2006). Immunoprecipitation (IP) was done using Myc trap agarose beads sourced from Chromotek (yta) in homogenization buffer (50 mM HEPES-KOH, pH 7.6; 1 mM EDTA; 140 mM KCl; 0.5% NP-40; 10% glycerol, 1× Protease Inhibitor Cocktail (Sigma 11836170001)). Post IP, the beads were rinsed thrice with homogenization buffer and boiled with Laemmli buffer. An equal amount of total protein (5 μg) in this lysate, assessed using a Pierce™ BCA Protein Assay Kit, was subjected to either an 8% (for whole UNC-104) or 12% (for fragments of UNC-104) denaturing PAGE and then transferred onto a supported nitrocellulose membrane. UNC-104 ubiquitin levels were assessed on blots with a normalized loading of UNC-104 protein amount assessed using the same lysate run on a previous blot probed for total UNC-104 intensity. The blots were probed with Ms anti-myc (AE010) (Lot no. 3500014031) (Dilution used 1:2500) to visualize the entire UNC-104 or UNC-104 fragments, Ms anti-ubiquitin (FK2) (Lot no. 3845622) (Dilution used 1:1000) to visualize ubiquitin, and Rb anti-actin (AC026) (Lot no. 9100026001) (Dilution used 1:5000) to visualize β-actin.

### Statistical analyses

Statistical analyses were performed using GraphPad Prism 9 or the SciPy stats module of Python 3.8 (Virtanen et al., 2020). The Shapiro–Wilk test was used to examine the normality of various distributions. Sample size estimates were calculated using G*Power 3.1.9.7 (Faul et al., 2009). Single comparisons between normally distributed data were performed using unpaired two-tailed Student’s *t*-test and those between non-normal distributions were performed using the Mann–Whitney U test. For multiple comparisons, one-way ANOVA with Dunnett’s multiple comparisons test was used for normal data, and Mann–Whitney–Wilcoxon test with Bonferroni multiple comparisons correction was used for non-normal data.

## Author Contributions

Conceptualization: V.S., S.P.K.; Methodology: V.S., S.P.K.; Theory: D.C., A.S., A.N.; Software: V.S., A.S., A.N., D.C.; Validation: V.S.; Formal analysis: V.S., A.S., A.N., D.C., S.P.K.; Investigation: V.S.; Resources: V.S., S.P.K., A.S., A.N., D.C., M.N.; Data curation: V.S., P.B., A.S.; Writing - original draft: V.S., S.P.K.; Theory portions of the paper is written by AS, AN and DC, Writing - review & editing: V.S., S.P.K., D.C., A.S., A.N., M.N.; Visualization: V.S., A.S.; Supervision: S.P.K.; Project administration: V.S., S.P.K.; Funding acquisition: S.P.K.

## List of supplementary legends and movies

**Supplementary Figure 1**

A) Schematic representing the regions imaged in the PLM neuron along with representative images showing FBXB-65::GFP intensity at the cell body, proximal process (70 µm from the cell body), the branch, and the distal end. Scale bar 10 µm.

B) Mean intensity of UNC-104::GFP (solid blue line) plotted with the standard deviation (shaded region) along the entire length of the PLM neuronal process in the strain TT2440 treated with either control, *uba-1* or *fbxb-65* RNAi (n>15). A linear fitting line (solid black line) to the intensity profile in the first 100 µm from the cell body extrapolated till the neuronal terminus to aid in visualization of the steep increase observed in the distal-most region. A residual plot shown below represents the mean intensity subtracted by the intensity expected from the fit at the specified region.

C) qPCR quantification of *unc-104* transcript levels using the ΔΔCt method in the strain TT2440 treated with either control, *uba-1* or *fbxb-65* RNAi. Results are plotted as Mean ± SEM (N=3 biological repeats); ns, non-significant (one-way ANOVA with Dunnett’s multiple comparisons test).

D) Intensity of UNC-104::GFP at the PLM synapses in the strain TT2440 treated with either control, *uba-1* or *fbxb-65* RNAi. Data are represented as violin plots with individual data points along with the median (dashed line), 25^th^ and 75^th^ percentile marked (dotted lines). *p<0.05 (one-way ANOVA with Dunnett’s multiple comparisons test).

**Supplementary Figure 2**

A) UNC-104::GFP intensity normalized to the average intensity of 5 frames pre-ablation plotted for a region including 5 µm of the cut site distal to the cell body in the strain TT2440 treated with either control, *uba-1* or *fbxb-65* RNAi. The solid line represents the mean with the shaded region representing the 95% confidence interval for ablation aligned (labeled as cut) time series intensities of n>15 for all conditions.

B) The fraction of UNC-104::GFP trajectories that have a run-length greater than 7 µm in a region 70 µm away from the cell body in the strain TT2440 treated with either control, *uba-1* or *fbxb-65* RNAi. Data are represented as violin plots with individual data points along with the median (dashed line), 25^th^ and 75^th^ percentile marked (dotted lines; ns, non-significant (one-way ANOVA with Dunnett’s multiple comparisons test).

C) Table of values of parameters A and µ found from fitting the equation (6) to the data in Fig 3G,H assuming γ=1 in the strain TT2440 treated with either control, *uba-1* or *fbxb-65* RNAi.

**Supplementary Figure 3**

A) Western blot analyses from anti-myc IP enriched fractions of TT3158 treated with either control, *uba-1, fbxb-65, uba-2, rfl-1, moc-3, uba-5* RNAi. The input served as a loading control probed with β-Actin. The two bands of UNC-104 PH::5xMYC are marked with arrow heads as HMW (High Molecular Weight) in red, LMW (Low Molecular Weight) in blue, and β-actin in black. Molecular weights (in kDa) marked with arrows.

B) Bar plot representing the mean intensity of the UNC-104 PH HMW normalized to the UNC-104 PH LMW from the western blot in A. Results are presented as Mean ± SD (N=3 biological repeats represented as filled circles). *p<0.05 (unpaired two-tailed *t*-test).

C) Western blot of worms expressing UNC-104 PH::5×MYC probed for anti-ubiquitin, stripped, and re-probed with anti-myc. Molecular weights (in kDa) marked with arrows.

D) Amino acid sequence of region deleted in the strain *unc-104(Δ1)*.

**Supplementary Figure 4**

A) Velocity of uninterrupted anterograde GFP::RAB-3 trajectories assessed from the strain TT2775 treated with either control or *fbxb-65* RNAi. Data are represented as violin plots with individual data points along with the median (dashed line), 25^th^ and 75^th^ percentile marked (dotted lines); ns, non-significant (Unpaired Student’s *t*-test two-tailed).

B) Fraction of anterograde GFP::RAB-3 trajectories with uninterrupted runs greater than 2 µm assessed from the strain TT2775 treated with either control or *fbxb-65* RNAi. Data represented as violin plots with individual data points along with the median (dashed line), 25^th^ and 75^th^ percentile marked (dotted lines). ***p<0.001 (Unpaired Student’s *t*-test two-tailed).

**Supplementary Figure 5**

A) Representative images of the PLM distal end in *xdKi3* (endogenous UNC-104::GFP) either in control or with *fbxb-65(0)*. Scale bar 10 µm.

B) Net displacement of anterogradely moving UNC-104::GFP trajectories assessed in *xdKi3* with either control or *fbxb-65(0)*. Data are represented as violin plots with individual data points along with the median (dashed line), 25^th^ and 75^th^ percentile marked (dotted lines); ns, non-significant (Unpaired Student’s *t*-test two-tailed).

C) Frequency distribution of background subtracted UNC-104 intensity averaged over an entire individual run assessed from the control *xdKi3*. The distribution is overlaid with a probability distribution function (red line) generated from our theoretical equation (6) assuming γ=1. Parameters were derived from the datasets by least square fitting.

D) Frequency distribution of background subtracted UNC-104 intensity averaged over an entire individual run assessed from *xdKi3* with the mutant *fbxb-65(0)*. The distribution is overlaid with a probability distribution function (red line) generated from our theoretical equation (6) assuming γ=1. Parameters were derived from the datasets by least square fitting.

E) Number of anterograde GFP::RAB-3 trajectories normalized to a 20 µm and 10 s region of the kymograph assessed in *xdKi3* with either control, *fbxb-65(0)* or *unc-104(Δ1)*. Data are represented as violin plots with individual data points along with the median (dashed line), 25^th^ and 75^th^ percentile marked (dotted lines). **p<0.01 ***p<0.001 (one-way ANOVA with Dunnett’s multiple comparisons test).

F) Velocity of uninterrupted anterograde GFP::RAB-3 trajectories assessed in either control, *fbxb-65(0)* or *unc-104(Δ1)*. Data are represented as violin plots with individual data points along with the median (dashed line), 25^th^ and 75^th^ percentile marked (dotted lines); ns, non-significant (one-way ANOVA with Dunnett’s multiple comparisons test).

G) Area integrated intensity of GFP::RAB-3 synapses assessed in either control, *fbxb-65(0)* or *unc-104(Δ1)*. Data are represented as violin plots with individual data points along with the median (dashed line), 25^th^ and 75^th^ percentile marked (dotted lines). ***p<0.001; ns, non-significant (one-way ANOVA with Dunnett’s multiple comparisons test).

**Movie 1: UNC-104::GFP in Control RNAi**

UNC-104::GFP in the PLM neuronal process 100 µm from the cell body. Imaged sequentially at 3 frames per second (fps), playback at 15 fps. Genotype: TT2440. Treatment: Control RNAi. Cell body on the right. Scale 10 µm.

**Movie 2: UNC-104::GFP in *fbxb-65* RNAi**

UNC-104::GFP in the PLM neuronal process 100 µm from the cell body. Imaged sequentially at 3 frames per second (fps), playback at 15 fps. Genotype: TT2440. Treatment: *fbxb-65* RNAi. Cell body on the right. Scale 10 µm.

**Movie 3: Knock-in UNC-104::GFP in wild type**

UNC-104::GFP in the PLM neuronal process 100 µm from the cell body. Imaged sequentially at 3 frames per second (fps), playback at 15 fps. Genotype: *xdKi3*. Cell body on the right. Scale 10 µm.

**Movie 4: Knock-in UNC-104::GFP in *fbxb-65(0)***

UNC-104::GFP in the PLM neuronal process 100 µm from the cell body. Imaged sequentially at 3 frames per second (fps), playback at 15 fps. Genotype: *xdKi3; fbxb-65(0)*. Cell body on the right. Scale 10 µm.

## Supporting information

All supplementary files

## Acknowledgments

We thank Mei Ding for the strain *xdKi3*, Kavya S. Pillai for building and characterizing the RNAi sensitive strain *sid-1(pk3321) him-5(e1490)*; *uIs71*; *jsIs1111*; *lin-15B(n744)*, Amal Mathew for building and characterizing the strain *sid-1(pk3321) him-5(e1490)*; *uIs71*; *jsIs821*; *lin-15B(n744)* and creating the *rab-3p*::UNC-104 construct. We thank Amruta Vasudevan for thoughtful discussions. Some strains were provided by the CGC, which is funded by the NIH Office of Research Infrastructure Programs (P40 OD010440). Research in the Sandhya Koushika’s lab is supported by grants from DAE (1303/2/2019/R&D-II/DAE/2079), PRISM (12-R&D-IMS-5.02-0202), and Howard Hughes Medical Institute International Early Career Scientist Grant 55007425. Debasish Chaudhuri acknowledges research grants from DAE (1603/2/2020/IoP/R&D-II/150288) and SERB, India (MTR/2019/000750), and thanks ICTS-TIFR, Bangalore, for an Associateship.

