## Supplementary material for "F-box protein FBXB-65 regulates anterograde transport of UNC-104 through modification near the PH domain": All supplementary files

### Supplementary Figure 1

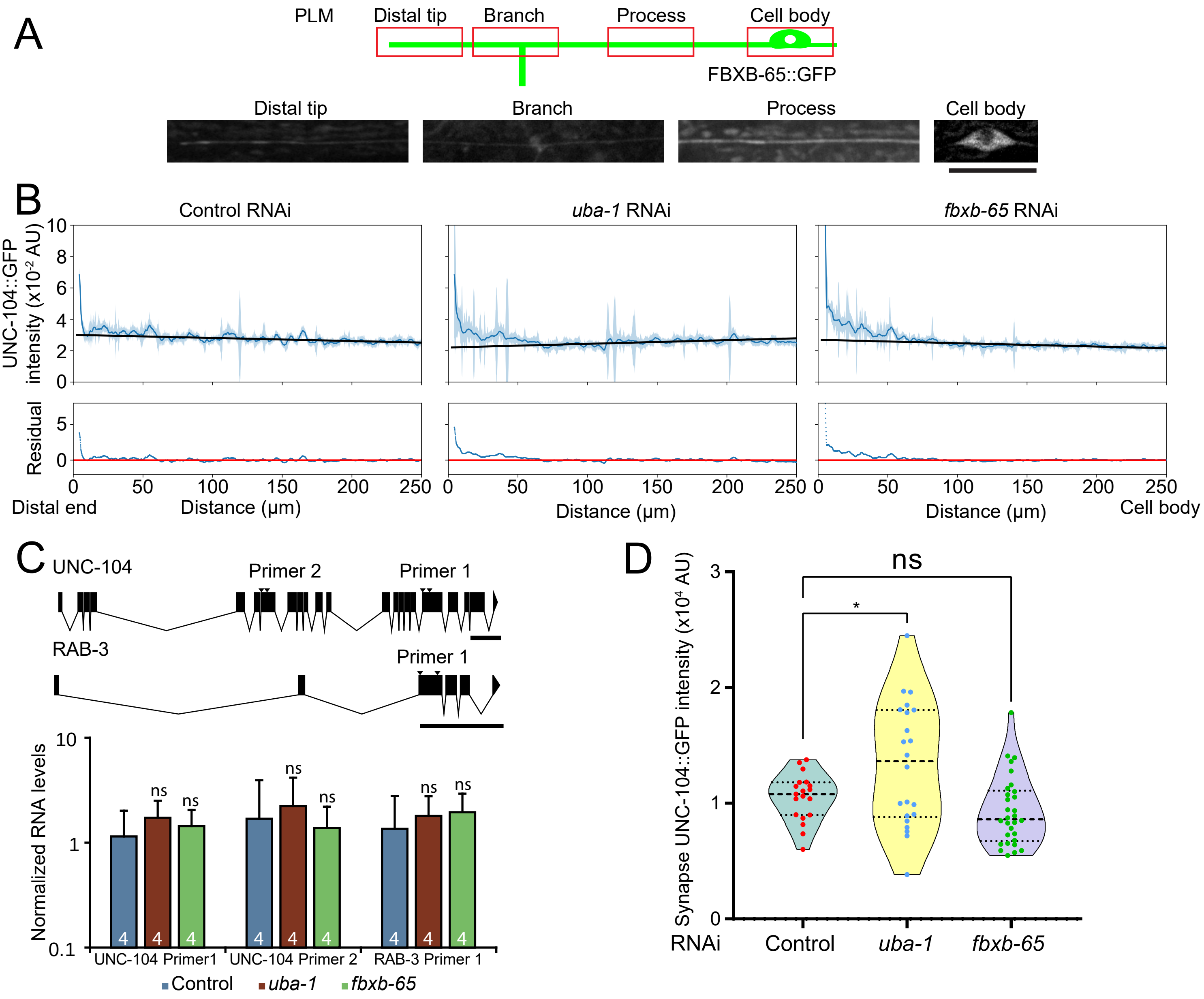

### Supplementary Figure 2

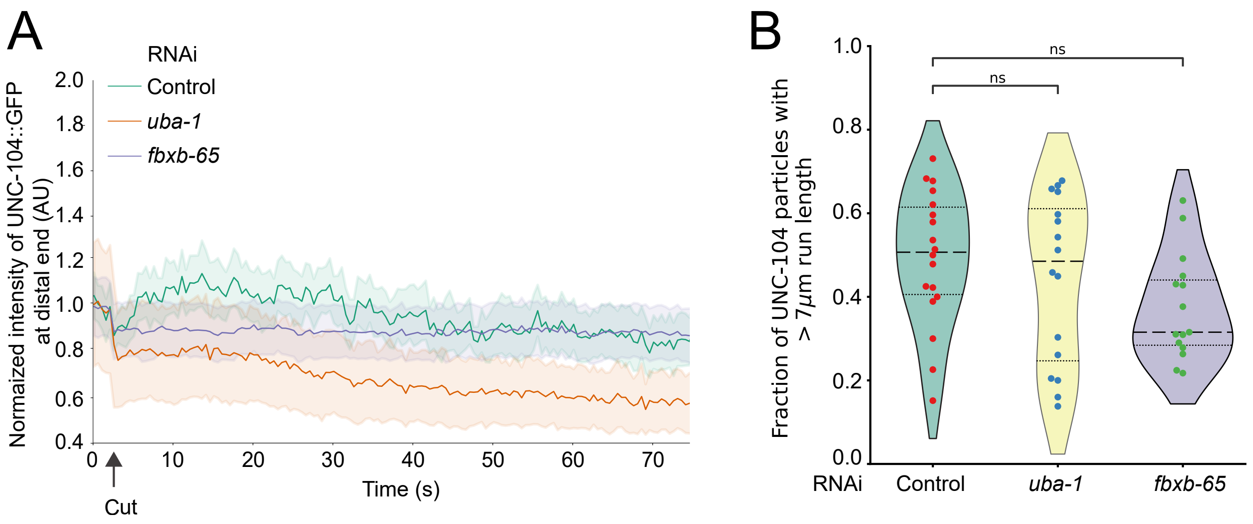

### Supplementary Figure 3

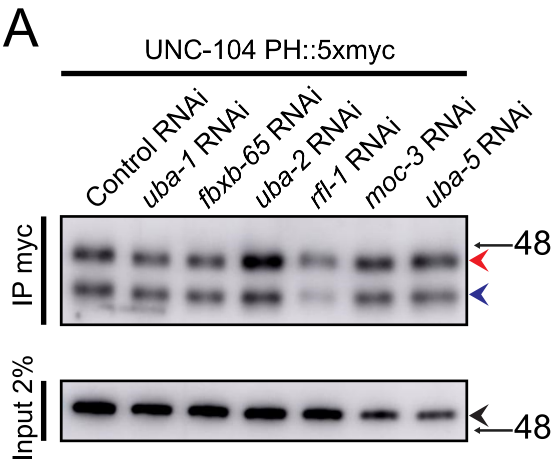


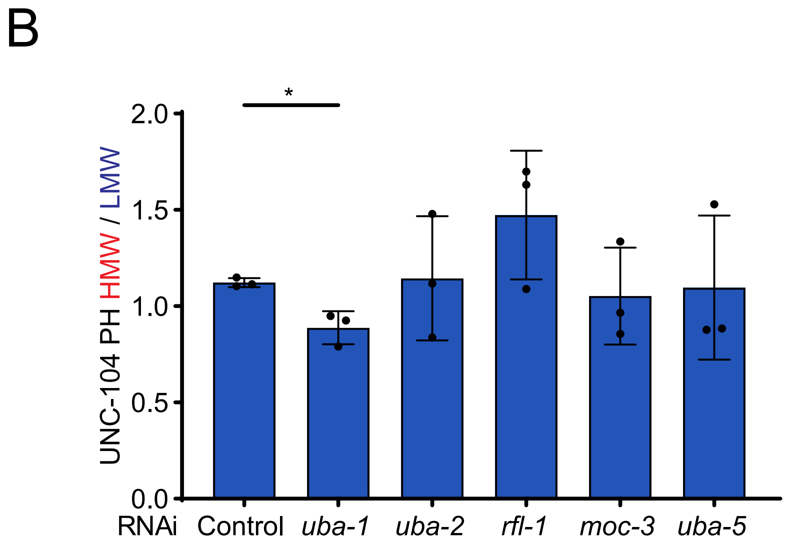


| RNAi | Control | *uba-1* | *uba-2* | *rfl-1* | *moc-3* | *uba-5* |
|---|---|---|---|---|---|---|
| Fold change compared to control | 1 | 0.79 | 1.01 | 1.31 | 0.93 | 0.98 |

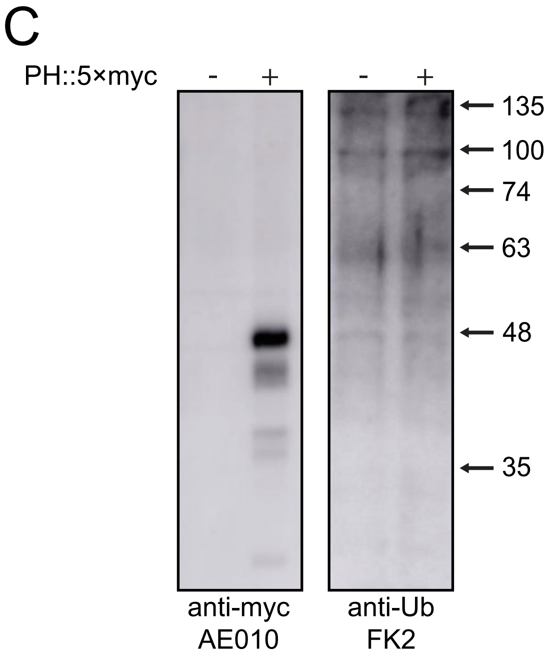


D

Δ1 PH sequence
PESIKLDEKDKGIVGKVLGLIRRKIPMNKDPPTGNK

### Supplementary Figure 4

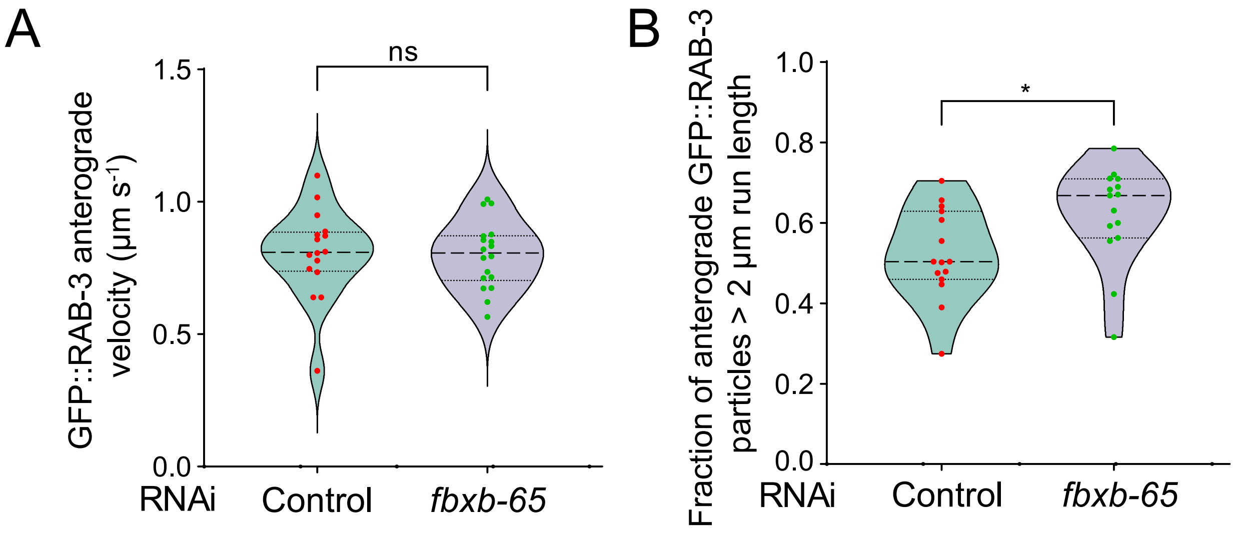

### Supplementary Figure 5

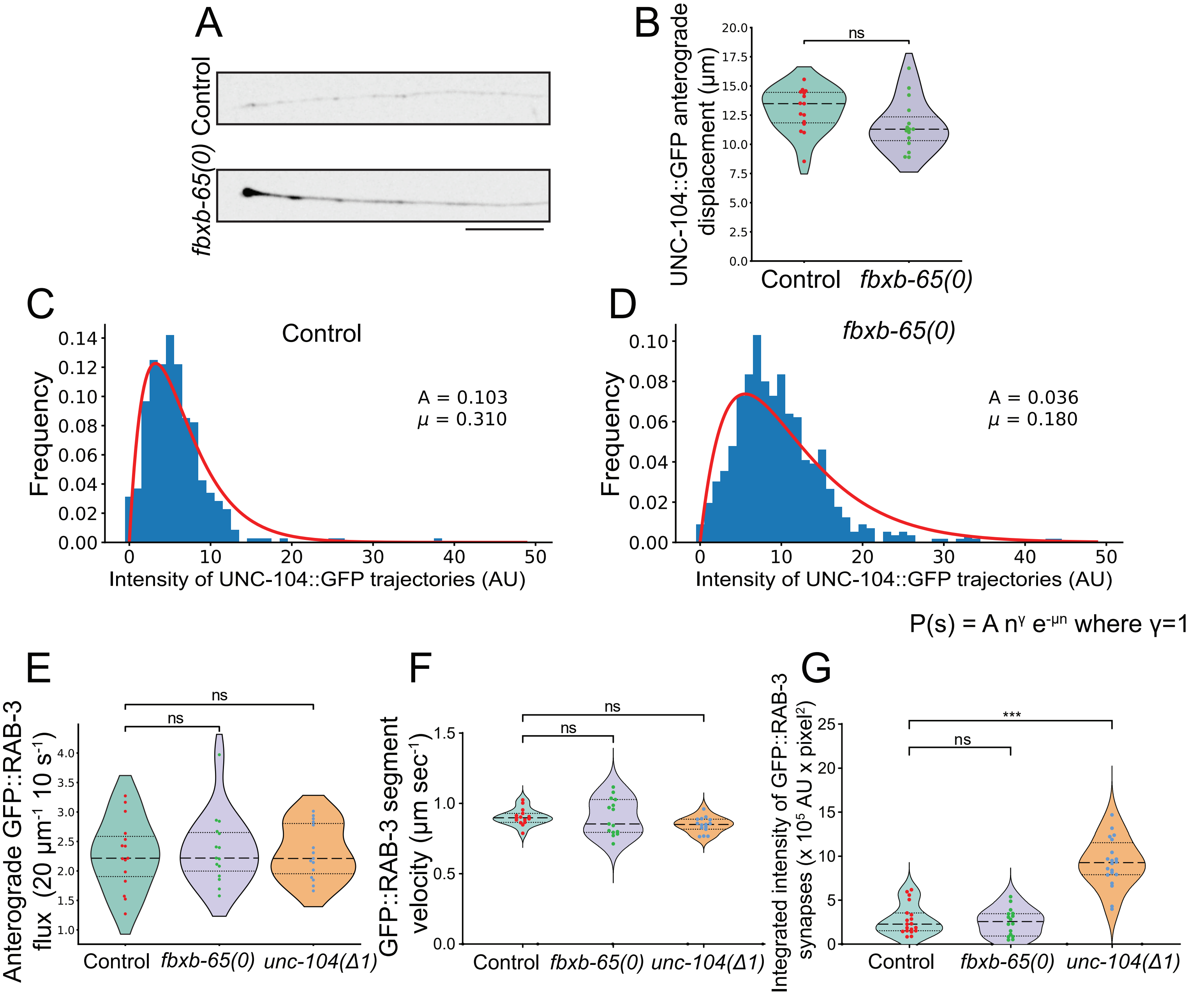

**Supplementary Table 1: List of strains**

| S. No. | Strain number | Genotype | Reference |
|---|---|---|---|
| 1 | N2 | Bristol wild type | (Brenner, 1974) |
| 2 | NM2689 | *jsIs821* [*mec-7p*::*gfp*::*rab-3*] | (Bounoutas et al., 2009) |
| 3 | TT385 | *unc-104(e1265tb120)* | (Kumar et al., 2010) |
| 4 | TU3401 | *sid-1(pk3321) uIs69* [(pCFJ90) *myo-2p*::*mCherry* + *unc-119p*::*sid-1*] V | (Calixto et al., 2010) |
| 5 | TU3568 | *sid-1(pk3321) him-5(e1490)* V; *lin-15B(n744)* X; *uIs71* [(pCFJ90) *myo-2p*::*mCherry* + *mec-18p*::*sid-1*] | (Calixto et al., 2010) |
| 6 | NM3764 | *jsIs1111 (mec-4p*::*unc-104*::*gfp*) | (Kumar et al., 2010) |
| 7 | XD5031 | *xdKi3(unc-104::gfp* knock-in*)* II | (Cong et al., 2021) |
| 8 | KG5148 | *unc-104*(*ce833*[5xMYC::AID::*unc-104+sup-1(e995)*]) | Stec et al (Unpublished) |
| 9 | NM2879 | *sam-4(js415) II; jsIs821 X* | (Zheng et al., 2014) |
| 10 | TT3591 | *tbEx525* [(pTT580) *myo-2p*::*mCherry* + (pTT731) *rab-3p*::UNC-104(1-390aa)::5xMYC] | This study |
| 11 | TT3554 | *tbEx515* [(pTT580) *myo-2p*::*mCherry* + (pTT732) *rab-3p*::UNC-104(380-874aa)::5xMYC] | This study |
| 12 | TT3536 | *tbEx502* [(pTT580) *myo-2p*::*mCherry* + (pTT733) *rab-3p*::UNC-104(845-1420aa)::5xMYC] | This study |
| 13 | TT3071 | *tbEx430* [(pTT540) *ttx-3p*::*gfp* + (pTT734) *rab-3p*::UNC-104(1386-1637aa)::5xMYC] | This study |
| 14 | TT3552 | *tbEx513* [(pTT580) *myo-2p*::*mCherry* + (pTT760) *rab-3p*::UNC-104(1386-1637aa del1386-1421)::5xMYC] | This study |
| 15 | TT3108 | *tbEx451* [(pTT579) *myo-3p*::*mCherry* + (pTT740) *rab-3p*::UNC-104(1386-1637aa del1422-1440)::5xMYC] | This study |
| 16 | TT3593 | *tbEx527* [(pTT580) *myo-2p*::*mCherry* + (pTT761) *rab-3p*::UNC-104(1386-1637aa del1441-1455)::5xMYC] | This study |
| 17 | TT3153 | *tbEx465* [(pTT580) *myo-2p*::*mCherry* + (pTT762) *rab-3p*::UNC-104(1386-1637aa del 1456-1476)::5xMYC] | This study |
| 18 | PHX7320 | *fbxb-65(syb7320)* | This study |
| 19 | PHX7293 | *unc-104b(syb7293)* (del1394-1430aa) | This study |
| 20 | TT3185 | *unc-104(ce833)* II; *sid-1(pk3321) uIs69* V | This study |
| 21 | TT3158 | *sid-1(pk3321) uIs69* V; *tbEx430* | This study |
| 22 | TT2440 | *sid-1(pk3321) him-5(e1490)* V; *uIs71*; *jsIs1111*; *lin-15B(n744)* X | This study |
| 23 | TT2775 | *sid-1(pk3321) him-5* V;*uIs71*;*lin-15b(n744) jsIs821* X | This study |
| 24 | TT3186 | *sam-4(js415)* II; *sid-1(pk3321) him-5(e1490)* V; *uIs71*; *lin-15B(n744) jsIs821* X | This study |
| 25 | TT3230 | *sam-4(js415)* II*;sid-1(pk3321) him-5(e1490)*; *uIs71*; *lin-15B(n744) jsIs821* X; *tbEx483*(*rab-3p*::UNC-104) | This study |
| 26 | TT3262 | *jsSi2013 tbSi520* [loxP mec-4Sp sng-1-L-mNG-C1 tbb-2 3' FRT3] IV | This study, generated from (Nonet 2023 preprint) |

**Supplementary Table 2: List of primers**

| S. No. | Primer number | Primer name | Primer sequence |
|---|---|---|---|
| 1 | TTpr1035 | RAB-3 Set1 qPCR | 5' GACTACATGTTCAAGctcctgataatcgg |
| 2 | TTpr1036 | | 5' gaatcccattgctccacgatagtagg |
| 3 | TTpr845 | UNC-104 Set1 qPCR | 5' ATGCACCAATTCAGAACAATAACGCATCTG |
| 4 | TTpr846 | | 5' CTCGAATACGTCCAACAACTAGTTCTTGAC |
| 5 | TTpr847 | UNC-104 Set2 qPCR | 5' CACTGAGTACTTTGAGATATGCCGATAGAGC |
| 6 | TTpr848 | | 5' GCTCATGTACATGAGCTGGCAATTTCG |
| 7 | TTpr40 | Actin qPCR | 5' GTAGACAATGGATCCGGAATGTGCAAGG |
| 8 | TTpr41 | | 5' GGTACTTGAGGGTAAGGATACCTCTCTTGG |
| 9 | TTpr1039 | UNC-104 Motor | 5' gatccccgggattggccATGTCATCGGTTAAAGTAGCTG |
| 10 | TTpr1040 | | 5' CAGACGTACAGGAGACACCCgagcaaaagcttatctctgag |
| 11 | TTpr1041 | UNC-104 Coiled coils | 5' gatccccgggattggccATGGGAATTGATGTCACAGACG |
| 12 | TTpr1042 | | 5' CCATGGTTTCGAATGGTTGGAgagcaaaagcttatctctgag |
| 13 | TTpr1043 | UNC-104 Stalk | 5' gatccccgggattggccATGTCGCCAGCTGATGGAG |
| 14 | TTpr1105 | | 5' GGATCCACCAACTGGAAACcaatttcagatggagcaaaagctta |
| 15 | TTpr1045 | UNC-104 PH | 5' gatccccgggattggccATGCCGGAAAGTATCAAGTTAGACG |
| 16 | TTpr1046 | | 5' GCTTTCTTGTACAAAGTGGTGgagcaaaagcttatctctgag |
| 17 | TTpr1047 | 5xMYC primer | 5' tctaggtaccTCATCCAGAACCTCCGAGGTCCTC |
| 18 | TTpr1138 | UNC-104 (Δ1) | 5' CCAAGCTTGGCCATGGCTCAAGAATTGAGTGATGAAAGTG |
| 19 | TTpr1139 | | 5' CAATTCTTGAGCCATGGCCAAGCTTGGGGATCCTCTAGAG |
| 20 | TTpr1102 | UNC-104 (Δ2) | 5' CTGGAAACAAAaaaagcttgatcaaatcctctcgatc |
| 21 | TTpr1103 | | 5' gatcaagcttttTTTGTTTCCAGTTGGTGGATCC |
| 22 | TTpr1140 | UNC-104 (Δ3) | 5' GATTTTGATTTCTGATCAGACACCGGTGATGTTATACTATTTG |
| 23 | TTpr1141 | | 5' TCACCGGTGTCTGATCAGAAATCAAAATCCGATCAGAACCTTG |
| 24 | TTpr1142 | UNC-104 (Δ4) | 5' CCACTCAAGCTTCGTTTCATTCGGCACAGGAGATCCGAAG |
| 25 | TTpr1143 | | 5' ATCTCCTGTGCCGAATGAAACGAAGCTTGAGTGGATCACG |
| 26 | TTpr1372 | *fbxb-65(syb7320)* screening | 5' aggcaccctatctagcagttgatgg |
| 27 | TTpr1373 | | 5' gctcctgaaaatctatgtttccaaaaacc |
| 28 | TTpr1375 | | 5' CCAAGTGGAGCTGGATACGATCAGATC |
| 29 | TTpr1322 | *unc-104(syb7293)* screening | 5' cgtcgacaacgtcgtgtacttgac |
| 30 | TTpr1318 | | 5' tcttgGTTGGTGGATCCTTGTTCATTGG |
| 31 | TTpr1142 | | 5' CCACTCAAGCTTCGTTTCATTCGGCACAGGAGATCCGAAG |

**Supplementary Table 3: List of plasmids**

| No. | Strain name | Details | Plasmid | Plasmid details | Source |
|---|---|---|---|---|---|
| 1 | *tbEx525* | Generated by injecting pTT731 (40 ng μl$^{-1}$) with myo-2p::gfp (50 ng μl$^{-1}$) as a co-injection marker | pTT731 | *rab-3p*::UNC-104(1-390aa)::5xMYC. Generated by 2 step in-fusion, first amplifying 5xMYC from the strain *unc-104(ce833)* using primers TTpr1040+1047, followed by using this product as a reverse primer and TTpr1139 as a forward primer to amplify the desired unc-104 fragment from the construct pSN8(*itr-1p*::*unc-104*::gfp). The final amplified product was inserted into the plasmid pHW393 (*rab-3p*:: GAL4-SK(DBD)::VP64::let-858 3'UTR) by replacing the *mCherry* insert using the XmaI and KpnI restriction sites | This study, pSN8(*itr-1p*::*unc-104*::*gfp*) was a gift from S. Niwa (Niwa 2016) , pHW393 (*rab-3p*:: GAL4-SK(DBD)::VP64::let-858 3'UTR) was a gift from Paul Sternberg (Addgene plasmid # 85583) |
| 2 | tbEx515 | Generated by injecting pTT732 (40 ng μl$^{-1}$) with *myo-2p::gfp* (50 ng μl$^{-1}$) as a co-injection marker | pTT732 | *rab-3p*::UNC-104(380-874aa)::5xMYC. Generated by 2 step in-fusion, first amplifying 5xMYC from the strain *unc-104(ce833)* using primers TTpr1042+1047, followed by using this product as a reverse primer and TTpr1141 as a forward primer to amplify the desired unc-104 fragment from the construct pSN8(*itr-1p*::*unc-104*::gfp). The final amplified product was inserted into the plasmid pHW393 (*rab-3p*:: GAL4-SK(DBD)::VP64::let-858 3'UTR) by replacing the *mCherry* insert using the XmaI and KpnI restriction sites | This study, pSN8(*itr-1p*::*unc-104*::*gfp*) was a gift from S. Niwa (Niwa 2016) , pHW393 (*rab-3p*:: GAL4-SK(DBD)::VP64::let-858 3'UTR) was a gift from Paul Sternberg (Addgene plasmid # 85583) |
| 3 | tbEx502 | Generated by injecting pTT733 (150 ng μl-1) with *myo-2p::gfp* (50 ng μl$^{-1}$) as a co-injection marker | pTT733 | *rab-3p*::UNC-104(845-1420aa)::5xMYC. Generated by 2 step in-fusion, first amplifying 5xMYC from the strain *unc-104(ce833)* using primers TTpr1044+1047, followed by using this product as a reverse primer and TTpr1143 as a forward primer to amplify the desired *unc-104* fragment from the construct pSN8(*itr-1p*::*unc-104*::gfp). The final amplified product was inserted into the plasmid pHW393 (*rab-3p*:: GAL4-SK(DBD)::VP64::let-858 3'UTR) by replacing the *mCherry* insert using the XmaI and KpnI restriction sites | This study, pSN8(*itr-1p*::*unc-104*::*gfp*) was a gift from S. Niwa (Niwa 2016) , pHW393 (*rab-3p*:: GAL4-SK(DBD)::VP64::let-858 3'UTR) was a gift from Paul Sternberg (Addgene plasmid # 85583) |
| 4 | tbEx430 | Generated by injecting pTT734 (40 ng μl-1) with *ttx-3*p::gfp (50 ng μl$^{-1}$) as a co-injection marker | pTT734 | *rab-3p*::UNC-104(1386-1637aa)::5xMYC. Generated by 2 step in-fusion, first amplifying 5xMYC from the strain *unc-104(ce833)* using primers TTpr1046+1047, followed by using this product as a reverse primer and TTpr1145 as a forward primer to amplify the desired *unc-104* fragment from the construct pSN8(*itr-1p*::*unc-104*::gfp). The final amplified product was inserted into the plasmid pHW393 (*rab-3p*:: | This study, pSN8(*itr-1p*::*unc-104*::*gfp*) was a gift from S. Niwa (Niwa 2016) , pHW393 (*rab-3p*:: GAL4-SK(DBD)::VP64::let-858 3'UTR) was a gift from Paul Sternberg (Addgene plasmid # 85583) |

| | | | | | |
|---|---|---|---|---|---|
| | | | | GAL4-SK(DBD)::VP64::let-858 3'UTR) by replacing the *mCherry* insert using the XmaI and KpnI restriction sites | |
| 5 | tbEx513 | Generated by injecting pTT760 (40 ng µl-1) with *myo-2p::gfp* (50 ng $\mu l^{-1}$) as a co-injection marker | pTT760 | *rab-3p*::UNC-104(1386-1637aa del1386-1421)::5xMYC. Generated by inverse PCR of the construct pTT734 deleting the region encoding (1386-1421aa) using the primers TTpr1138 and TTpr1139. A 27 bp overlap was maintained in these primers to aid in *in vivo* (*E. coli* DH5a) DNA assembly | This study. |
| 6 | tbEx451 | Generated by injecting pTT740 (40 ng µl-1) with *myo-3p::gfp* (50 ng $\mu l^{-1}$) as a co-injection marker | pTT740 | rab-3p::UNC-104(1386-1637aa del1422-1440)::5xMYC Generated by inverse PCR of the construct pTT734 deleting the region encoding (1422-1440aa) using the primers TTpr1102 and TTpr1103. A 23 bp overlap was maintained in these primers to aid in *in vivo* (*E. coli* DH5a) DNA assembly | This study. |
| 7 | tbEx527 | Generated by injecting pTT761 (40 ng µl-1) with *myo-2p::gfp* (50 ng $\mu l^{-1}$) as a co-injection marker | pTT761 | *rab-3p*::UNC-104(1386-1637aa del1441-1455)::5xMYC. Generated by inverse PCR of the construct pTT734 deleting the region encoding (1441-1455aa) using the primers TTpr1140 and TTpr1141. A 29 bp overlap was maintained in these primers to aid in *in vivo* (*E. coli* DH5a) DNA assembly | This study. |
| 8 | tbEx465 | Generated by injecting pTT762 (40 ng µl-1) with *myo-2p::gfp* (50 ng $\mu l^{-1}$) as a co-injection marker | pTT762 | *rab-3p*::UNC-104(1386-1637aa del 1456-1476)::5xMYC. Generated by inverse PCR of the construct pTT734 deleting the region encoding (1456-1476aa) using the primers TTpr1142 and TTpr1143. A 34 bp overlap was maintained in these primers to aid in *in vivo* (*E. coli* DH5a) DNA assembly | This study. |

**Supplementary Table 4: Associated with Fig. 1**

| Figure | Test | Comparison | p-value |
|---|---|---|---|
| 1C | One-way ANOVA with Dunnett's multiple comparisons test | Control vs. *uba-1* | 0.0103 |
| | | Control vs. *fbxb-65* | 0.0881 |
| | | *uba-1* vs *fbxb-65* | 0.2384 |
| 1E | One-way ANOVA with Dunnett's multiple comparisons test | Control vs. *uba-1* | 0.0410 |
| | | Control vs. *fbxb-65* | 0.9980 |
| | | *uba-1* vs *fbxb-65* | 0.0381 |

**Supplementary Table 5: Associated with Fig. 2**

| Figure | Test | Comparison | p-value |
|---|---|---|---|
| 2B | Unpaired Student's *t*-test two-tailed | Control vs. *fbxb-65* | 0.0921 |
| 2C | Unpaired Student's *t*-test two-tailed | Control vs. *fbxb-65* | 0.1763 |
| 2D | Unpaired Student's *t*-test two-tailed | Control vs. *fbxb-65* | 0.0106 |
| 2E | Mann–Whitney test two-tailed | Control vs. *fbxb-65* | 0.0018 |
| 2F | Mann–Whitney test two-tailed | Control vs. *fbxb-65* | 0.0298 |

**Supplementary Table 6: Associated with Fig. 3**

| Figure | Test | Comparison | p-value |
|---|---|---|---|
| 3D | Mann–Whitney–Wilcoxon test with Bonferroni correction | Control vs. *uba-1* | 0.04143 |
| | | Control vs. *fbxb-65* | 0.03136 |
| 3E | Mann–Whitney–Wilcoxon test with Bonferroni correction | Control vs. *uba-1* | 0.00285 |
| | | Control vs. *fbxb-65* | 0.01922 |
| 3G | Mann–Whitney–Wilcoxon test with Bonferroni correction | Control vs. *uba-1* | 0.06268 |
| | | Control vs. *fbxb-65* | 0.02134 |
| 3H | Mann–Whitney–Wilcoxon test with Bonferroni correction | Control vs. *uba-1* | 0.04753 |
| | | Control vs. *fbxb-65* | 0.01919 |

**Supplementary Table 7: Associated with Fig. 4**

| Figure | Test | Comparison | p-value |
|---|---|---|---|
| 4D | Unpaired Student's *t*-test two-tailed | Control vs. *fbxb-65* | 0.0158 |

**Supplementary Table 8: Associated with Fig. 5**

| Figure | Test | Comparison | p-value |
|---|---|---|---|
| 5B | Unpaired Student's *t*-test two-tailed | Control vs. *fbxb-65* | 0.0412 |
| 5C | Unpaired Student's *t*-test two-tailed | Control vs. *fbxb-65* | 0.3178 |
| 5D | Unpaired Student's *t*-test two-tailed | Control vs. *fbxb-65* | 0.0031 |
| 5F | Unpaired Student's *t*-test two-tailed | Control vs. *fbxb-65* | 0.0019 |
| 5G | Unpaired Student's *t*-test two-tailed | Control vs. *fbxb-65* | <0.0001 |

**Supplementary Table 9: Associated with Fig. 6**

| Figure | Test | Comparison | p-value |
|---|---|---|---|
| 6C | Unpaired Student's *t*-test two-tailed | Control vs. *fbxb-65* | 0.0012 |
| 6D | Unpaired Student's *t*-test two-tailed | Control vs. *fbxb-65* | 0.0002 |
| 6G | One-way ANOVA with Dunnett's multiple comparisons test | Control vs. *fbxb-65(Δ)* | 0.4828 |
| | | Control vs. *unc-104(PHΔ1)* | 0.042 |
| 6H | One-way ANOVA with Dunnett's multiple comparisons test | Control vs. *fbxb-65(Δ)* | <0.0001 |
| | | Control vs. *unc-104(PHΔ1)* | 0.0034 |

**Supplementary Table 10: Associated with Fig. 7**

| Figure | Test | Comparison | p-value |
|---|---|---|---|
| 7B | Unpaired Student's *t*-test two-tailed | Control vs. *fbxb-65(Δ)* | 0.0034 |
| 7D | One-way ANOVA with Dunnett's multiple comparisons test | Control vs. *fbxb-65* | 0.0457 |
| | | Control vs. UNC-104 O/E | 0.0013 |
| 7G | One-way ANOVA with Dunnett's multiple comparisons test | Control vs. *fbxb-65(0)* | 0.0069 |
| | | Control vs. *unc-104(PHΔ1)* | <0.0001 |
| 7H | One-way ANOVA with Dunnett's multiple comparisons test | Control vs. *fbxb-65(0)* | 0.8651 |
| | | Control vs. *unc-104(PHΔ1)* | <0.0001 |

**Supplementary Table 11: Associated with Fig. 8**

| Figure | Test | Comparison | p-value |
|---|---|---|---|
| 8B | One-way ANOVA with Dunnett's multiple comparisons test | Control vs. *fbxb-65(0)* | <0.0001 |
| | | Control vs. *unc-104(PHΔ1)* | <0.0001 |
| 8C | Kruskal-Wallis with Dunn's multiple comparisons test | Control vs. *fbxb-65(0)* | >0.9999 |
| | | Control vs. *unc-104(PHΔ1)* | 0.5465 |
| 8D | One-way ANOVA with Dunnett's multiple comparisons test | Control vs. *fbxb-65(0)* | <0.0001 |
| | | Control vs. *unc-104(PHΔ1)* | 0.0021 |

**Supplementary Table 12: Associated with Fig. S1**

| Figure | Test | Comparison | p-value |
|---|---|---|---|
| Supp Fig 1C | One-way ANOVA with Dunnett's multiple comparisons test | Control vs. *uba-1* UNC-104 Set 1 | 0.4819 |
| | | Control vs. *fbxb-65* UNC-104 Set 1 | 0.8193 |
| | One-way ANOVA with Dunnett's multiple comparisons test | Control vs. *uba-1* UNC-104 Set 2 | 0.8796 |
| | | Control vs. *fbxb-65* UNC-104 Set 2 | 0.9573 |
| | One-way ANOVA with Dunnett's multiple comparisons test | Control vs. *uba-1* RAB-3 Set 1 | 0.8170 |
| | | Control vs. *fbxb-65* RAB-3 Set 1 | 0.6995 |
| Supp Fig 1D | One-way ANOVA with Dunnett's multiple comparisons test | Control vs. *uba-1* | 0.0313 |
| | | Control vs. *fbxb-65* | 0.4369 |

**Supplementary Table 13: Associated with Fig. S2**

| Figure | Test | Comparison | p-value |
|---|---|---|---|
| Supp Fig 2B | One-way ANOVA with Dunnett's multiple comparisons test | Control vs. *fbxb-65* | 0.0807 |
| | | Control vs. *uba-1* | 0.5677 |

**Supplementary Table 14: Associated with Fig. S3**

| Figure | Test | Comparison | p-value |
|---|---|---|---|
| Supp Fig 3B | Unpaired Student's *t*-test two-tailed | Control vs. *uba-1* | 0.0103 |
| | Unpaired Student's *t*-test two-tailed | Control vs. *uba-2* | 0.9096 |
| | Unpaired Student's *t*-test two-tailed | Control vs. *rfl-1* | 0.1440 |
| | Unpaired Student's *t*-test two-tailed | Control vs. *moc-3* | 0.6582 |
| | Unpaired Student's *t*-test two-tailed | Control vs. *uba-5* | 0.9101 |

**Supplementary Table 15: Associated with Fig. S4**

| Figure | Test | Comparison | p-value |
|---|---|---|---|
| Supp Fig 4A | Unpaired Student's *t*-test two-tailed | Control vs. *fbxb-65* | 0.9107 |
| Supp Fig 4B | Unpaired Student's *t*-test two-tailed | Control vs. *fbxb-65* | <0.0001 |
| Supp Fig 4D | Mann–Whitney test two-tailed | Control vs. *fbxb-65* | 0.0607 |
| Supp Fig 4E | Unpaired Student's *t*-test two-tailed | Control vs. *fbxb-65* | 0.0015 |

**Supplementary Table 16: Associated with Fig. S5**

| Figure | Test | Comparison | p-value |
|---|---|---|---|
| Supp Fig 5B | Unpaired Student's *t*-test two-tailed | Control vs. *fbxb-65* | 0.0825 |
| Supp Fig 5E | One-way ANOVA with Dunnett's multiple comparisons test | Control vs. *fbxb-65(0)* | 0.8960 |
| | | Control vs. *unc-104(Δ1PH)* | 0.9739 |
| Supp Fig 5F | One-way ANOVA with Dunnett's multiple comparisons test | Control vs. *fbxb-65(0)* | 0.9992 |
| | | Control vs. *unc-104(Δ1PH)* | 0.1540 |
| Supp Fig 5G | One-way ANOVA with Dunnett's multiple comparisons test | Control vs. *fbxb-65(0)* | 0.8985 |
| | | Control vs. *unc-104(Δ1PH)* | <0.0001 |

**Supplementary Table 17: Associated with Fig. 3G,H,** Estimates of A and μ

| | Control RNAi | *fbxb-65* RNAi |
|---|---|---|
| A | $5.00 \times 10^{-5}$ | $9.06 \times 10^{-6}$ |
| μ | $7.33 \times 10^{-3}$ | $3.05 \times 10^{-3}$ |
